# Charting the single cell transcriptional landscape governing visual imprinting

**DOI:** 10.1101/2025.06.23.660422

**Authors:** Vincenzo Lagani, Lela Chitadze, Ana Cecilia Gonzalez Alvarez, Teimuraz Bokuchava, Giulia Sansone, Veriko Bokuchava, Lia Tsverava, Aragorn Jones, Steven Lisgo, Xabier Martinez De Morentin, Robert Lehmann, Leena Ali Ibrahim, Jesper Tegner, Brian J. McCabe, Zaza Khuchua, David Gomez Cabrero, Revaz Solomonia

## Abstract

Memory-related transcriptional events in brain remain poorly understood. Visual imprinting is a form of learning in which young animals develop preferences through early exposure to specific stimuli. In chicks, visual imprinting memory is stored in the intermediate medial mesopallium (IMM) of the forebrain. To investigate learning-associated molecular changes, we performed single-nucleus RNA sequencing of the left IMM in strongly imprinted chicks and untrained controls. This analysis generated the first classification of cells composing the IMM, identifying as a result over 30 cell clusters with distinct transcriptional differences putatively linked to memory formation, nearly half of them in long non-coding RNAs (lncRNAs). Follow-up analysis on selected genes, confirmed that the gene expression levels of two lncRNAs and protein levels of FOXP2, RORA (transcription factors), LUC7L (splicing factor), and ROBO1 (axon guidance molecule) correlate with memory strength, reflecting either innate learning potential or imprinting experience. Additionally, among the confirmed lncRNAs, the brain- and avian-specific lncRNA ENSGALG00010007489 is enriched in the nuclei of specific glutamatergic clusters and its association with imprinting was further confirmed through quantitative multi-probe in situ hybridization. These findings offer the first single-cell resolution map of transcriptional changes underlying memory formation in the avian brain.

## Introduction

Memory is the capacity to retain learned information and is of fundamental importance to animals. While different types of memory have distinct operational characteristics, they share certain molecular mechanisms, particularly those subserving synaptic plasticity^1^. Study of a given type of memory may therefore elucidate not only the specific molecular details of that particular process, but also extend understanding of generally applicable molecular mechanisms of memory^2^. The specific gene expression programs activated in distinct brain cell subtypes during memory encoding, consolidation, and retrieval remain poorly characterized, particularly at single-cell resolution. Understanding the neural mechanisms of learning and memory is crucial for both scientific advancement and practical applications. In particular, the rational treatment of mental illness, especially the amnesias, requires detailed knowledge of how the brain stores information^3^.

A particularly suitable type of learning for memory studies is visual imprinting in chicks. Visual imprinting is a form of learning that occurs early in life, in which the young animal rapidly forms a strong and lasting attachment to a visual stimulus after brief exposure to it^3–7^. Visual imprinting is characterized by highly specific recognition memory, making it a powerful tool for studying the biological basis of memory. In domestic chicks, this form of learning offers key advantages for memory research: (i) imprinting may be studied soon after hatching when chicks do not require food, ensuring standardized nutritional conditions; (ii) the sensory environment during incubation and rearing can be tightly controlled, minimizing extraneous sensory input and allowing for clearer detection of learning-related neural changes; and (iii) newly hatched chicks have a diverse behavioral repertoire, enabling precise behavioral assessments of learning. A commonly used measure of imprinting strength, and thus memory, is the preference score, which quantifies the chick’s approach toward the training stimulus relative to a novel, unfamiliar stimulus^3,6,7^.

Beyond these operational advantages, a specific brain region has been identified that plays a critical role in the learning and memory processes of chick visual imprinting ^3,6,8^. This region is the *intermediate medial mesopallium* (IMM), previously known as the *intermediate and medial hyperstriatum ventrale* (IMHV)^9^, which has been identified as a site of memory of the imprinting stimulus. Using regression analysis, it is possible to assess whether molecular changes in the IMM (and other brain regions) are attributable to memory formation consequent on learning, rather than to side effects of the training procedure. Criteria for identifying learning-related changes using linear modeling and variance partitioning have been documented^10^. Variance that is unexplained by covariation with learning (i.e., residual variance from a regression with preference score) provides information enabling discrimination between a neural change resulting from learning and learning ability existing before training (a predisposition to learn)^10^.

Visual imprinting in chicks is associated with hemispheric asymmetry at behavioral, molecular, and electrophysiological levels^3,6,7,11^. Learning-related molecular changes are typically more strongly expressed in the left IMM, especially 24 h after training^3,6,7,10,12–17^. For this reason, the present study focuses on the left IMM.

We have previously demonstrated a progression of learning-related molecular changes in the IMM over the 24 h after training^7^. At least some of these changes are not uniformly distributed among IMM cells and occur at different times after training^18–22^. For example, learning-related up-regulation of Fos protein in parvalbumin-containing, but not calbindin-containing, GABA neurons occurs 1 h after the end of training in the IMM but not subsequently^22^. In contrast, a learning-related change in NMDA receptor binding occurs several hours later^18,19^. To further understand the molecular mechanisms of memory, it is important to know which molecular changes are occurring and in which cell types they take place.

Single cells/nuclei omics technologies have revolutionized molecular biology, allowing investigation of the heterogeneity of living tissues at the cellular level^23^. Publicly available atlases^24,25^ of brain cell transcriptomics profiles, together with appropriate technology, enable detailed study of neural plasticity and memory at the cellular level^26,27^, increasing understanding of fundamental mechanisms and thus establishing a rational basis for the successful treatment of neurological disorders^28,29^. No previous study has investigated imprinting-associated molecular changes in the chick IMM with single-cell RNA-seq technology.

In the present study, we compared single-nucleus transcriptome profiles of the left IMM between untrained chicks and good learners – those that achieved high preference scores during training (see Methods). We collected tissue 24 h after the end of the training, a time point when numerous long-term, learning-dependent changes occur in the IMM^7,10,12–17,30^. For the first time, our findings have enabled the identification of cell-type-specific gene expression differences - both coding and non-coding - between good learners and untrained chicks.

Further analyses have revealed that for one long non-coding RNA (lncRNA), ENSGALG00010007489 (here identified as GLUBK89) at the transcript level, and three protein coding genes – LUC7L, FOXP2, and RORA at the protein level – display strong correlations between preference score and expression level, indicating changes attributable to learning during training. In contrast, changes in lncRNA ENSGALG00010026609 (lncRNA6609) and ROBO1 likely reflect learning capacity, suggesting a predisposition to learn existing prior to training. These validation experiments were performed in four brain regions (left and right IMM, left and right posterior pole of neostriatum, PPN, a control brain region not involved in imprinting). Moreover, the results indicate that the magnitude of learning-related gene-expression changes within specific cellular clusters is sufficiently large to be detected at the whole-tissue level.

Finally, fluorescence *in situ* hybridization (FISH) experiments confirmed that GLUBK89 is localized in the nuclei of putative glutamatergic neurons in the IMM. Expression of GLUBK89 in these neurons was significantly higher in good learner chicks compared to untrained controls, with poor learners exhibiting intermediate expression levels. These findings indicate that this lncRNA contributes to memory processing within a subset of glutamatergic neurons.

Our results provide insight into memory mechanisms of imprinting and are potentially of general importance in understanding the molecular basis of memory.

## Results

### IMM cellular composition suggests homology with deep layers of the mammalian association cortex

We followed an established protocol for training and testing^21^ (see Methods) to assess imprinting behavior. In short, 24 hours after hatching, chicks were placed in a running wheel and exposed to a training stimulus consisting of a cuboid rotating red light (Figure 1a). Ten minutes after training, a preference test was conducted to measure memory strength. Chicks with preference scores above 80 were considered good learners, having mostly approached the original training stimulus rather than the alternative one (a rotating blue light, Figure 1a). Twenty-four hours after the preference test, the chicks were decapitated, and the left IMM removed. Untrained chicks, which were never exposed to the imprinting stimulus (Figure 1a), served as controls.

**Figure 1:**
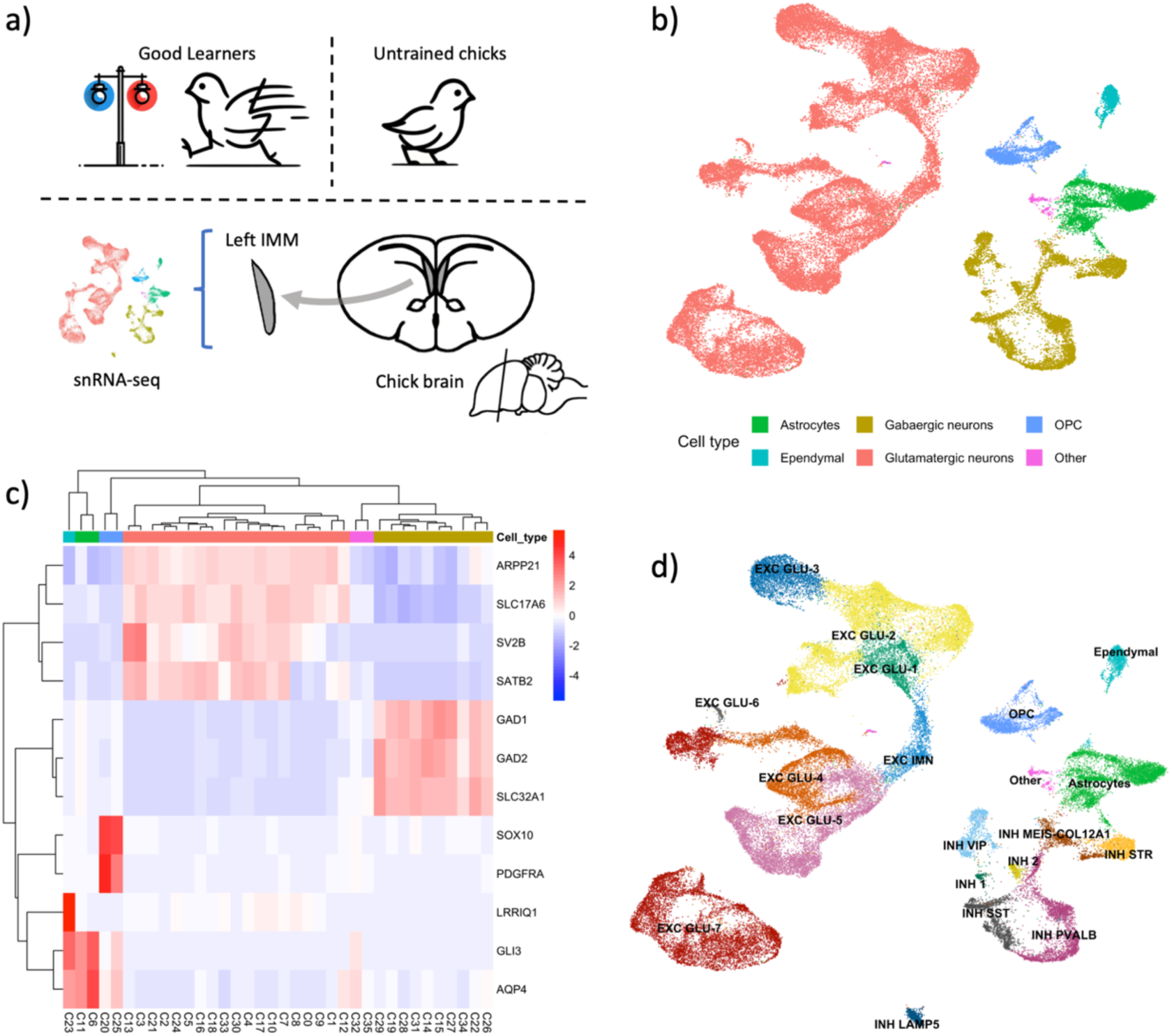
(a) Graphical presentation of the study design. 22-28h old chicks are either subjected to a preference test for assessing their imprinting or left untrained. Trained chicks are selected only if consistently directing themselves towards the training stimulus (good learners; see methods for details). The left portion of the intermediate and medial mesopallium (IMM) is collected from all selected chicks. Single nuclei transcriptomics data are then measured. (b) UMAP visualizing the 54763 IMM cells profiled with snRNA-seq. Color indicates major cell classifications: astrocytes, oligodendrocyte precursors (OPC), as well as GABAergic and glutamatergic neurons. (c) Heatmap showing the expression levels of gene markers for major cell types. (d) Localization of cell subtypes, see main text for details.

We processed left IMM samples from a total of 7 chicks (3 good learners and 4 untrained chicks, Supplementary Figure S1) using single-nucleus RNA-seq technology (see Methods), obtaining transcriptomic profiles for a total of 54,763 nuclei, which were grouped into 36 clusters (Supplementary Table S1 and Supplementary Figure S2).

We contrasted our snRNA-seq data with a recently published single-cell chicken brain scRNA-seq atlas^31^, to annotate our clusters on the basis of its classifications (see Methods and Supplementary Table S2). Major cell-types could be identified confidently, as confirmed by well-established marker genes^32–34^ (Figure 1b,c): GABAergic neurons (n=8,999; GAD1, GAD2, SLC32A1), glutamatergic neurons (n=37,582; ARPP21, SLC17A6, SV2B, SATB2), astrocytes (n=4,472; AQP4, GLI3), and oligodendrocyte precursors (OPC; n=2,251; PDGFRA and SOX10).

Beyond these major cell types, some clusters were not readily identified by this analysis. Cluster 32 (C32) expresses markers characteristic of vasculature (n=284; INPP5D, FLT1, and ADGRL4) while C35 is characterized by markers of the immune system (n=76; B2M, GPX1, SLC4A1). Cluster C23 (n = 1,099) presents a distinct marker signature, including the LRRIQ1, LRGUK and NEK10 genes, which are associated with “cilium movement involved in cell motility” and related gene ontology biological processes (Supplementary Table S3). We hypothesize that this group corresponds to ependymal cells.

With the overall cellular composition of the IMM defined, we next focused on finer-grained classification. Recent works^31,35^ have highlighted that GABAergic inhibitory neurons are largely conserved across amniotes. In contrast, no clear unequivocal homology has been established yet between avian and mammalian excitatory neuron subtypes.

Consistent with these observations, label transfer identifies known GABAergic subtypes in our data (Supplementary Table S2). LAMP5 inhibitory neurons form a distinct cluster, clearly separate from the other cells (Figure 1d). Other notable inhibitory subtypes include VIP- and MEIS-expressing neurons, as well as striatal-like inhibitory neurons (STR)^36^ and somatostatin (SST) cells^37^ (Figure 1d). Expression of key markers provides additional evidence for this classification (Supplementary Figures S3 – S14).

We next examined the organization of glutamatergic neurons. Our glutamatergic clusters present a high degree of transcriptional similarity with the excitatory subgroups identified in the chicken brain single-cell atlas (see Methods and Supplementary Table S2), and we therefore adopt the corresponding classification. Notably, a substantial fraction of glutamatergic neurons maps to subtypes resembling excitatory neurons of the deep layers of the mammalian cortex. Specifically, 9,269 (24.66%) neurons are classified as GLU-7, whose transcriptomics are most similar to layer 6 (L6) excitatory neurons, while 8,495 (22.60%) cells are classified as GLU-2, corresponding most closely to layer 4 and layer 5 (L4/L5) cortical glutamatergic neurons.

Finally, we applied self-assembling manifold mapping^38^ (SAMap), an algorithm that exploits similarities between protein sequences to match scRNA-seq profiles across species (Methods), as a confirmatory analysis of our findings. We used a single-cell transcriptomics atlas of the mouse cortex and hippocampus as a reference^32^. This analysis confirmed several results obtained through label transfer from the chicken brain cell atlas (Supplementary Table S2 and S4).

In summary, we annotated cells at two different resolutions: at a coarser level, we confidently identified three major cell types (glutamatergic neurons, GABAergic neurons, and non-neuronal cells), each spanning multiple clusters. At a finer level, we identified 21 cell subtypes, as summarized in Supplementary Table S2.

Together, these results provide the first detailed classification of the transcriptomic profiles of cells composing the IMM, a brain region strongly implicated in imprinting and passive avoidance memory in chicks^3^.

### Good learners and untrained chicks exhibit distinct transcriptomic profiles

We first examined cell-type representation in good learners and untrained chicks, to disentangle transcriptional changes associated with learning from potential differences in cellular composition.

Cells from both experimental groups were detected across all annotated subtypes (Figure 2a). Major cell types were robustly represented across all samples and did not exhibit significant differences in cell proportions between good learners and untrained chicks (Supplementary Figure S15).

**Figure 2:**
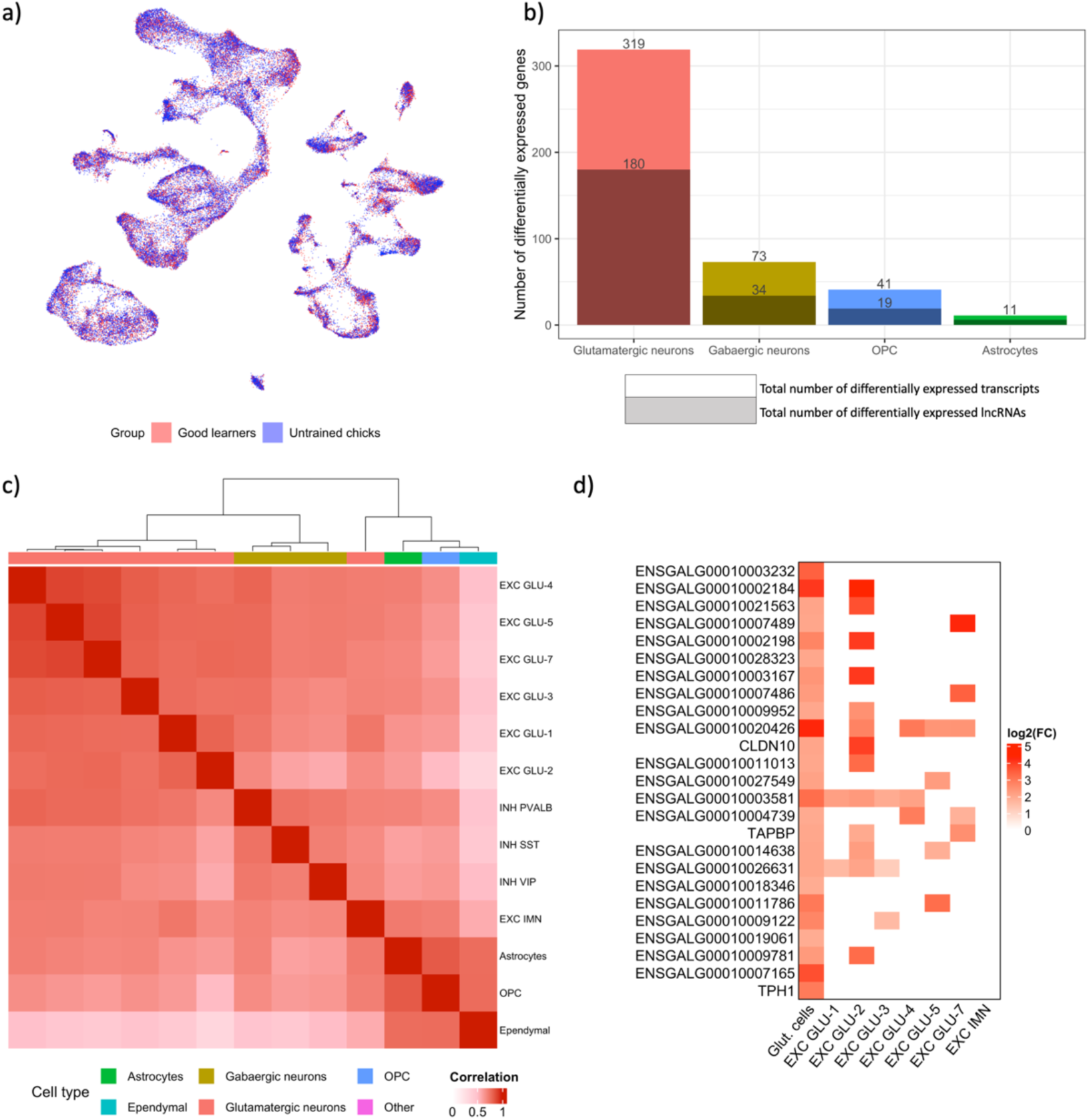
(a) UMAP visualizing IMM cells according to the group of their sample of origin, red good learners and blue untrained chicks. (b) Number of statistically significant transcriptomic changes (log FC > 1 in absolute value, FDR <= 0.05). Darker shaded bars indicate the number of non-coding RNA found differentially expressed. Six lncRNAs were differentially expressed for Astrocytes (c) Correlations among log2 fold changes across different clusters represented as heatmap. The dendrogram on top of the heatmap mostly splits clusters according to their original cell type, i.e., glutamatergic and GABAergic neurons, as well as non-neuronal cells. (d) Log2 FC for the transcripts with the highest increase of expression in glutamatergic cells. The log2 FC is reported for results with FDR <= 0.05, as well as with at least 50 cells in each pseudo-bulk sample (see Methods for details).

In contrast, eight cell subtypes contained fewer than 50 cells in at least one of the seven samples: EXC GLU-6, INH 1, INH 2, INH LAMP5, INH STR, Blood and Immune System, Microglia/Endothelial/Vascular, and INH MEIS2/COL12A1. We therefore refrained from drawing conclusions for these subtypes, either with respect to cell proportion changes or differential gene expression, as differences cannot be reliably interpreted in the presence of low per-sample cell counts^39^. The remaining subtypes showed no significant differences in overall cell abundance between experimental groups (Supplementary Figure S15).

Next, we performed a pseudo-bulk differential expression analysis between conditions (see Methods), both for major cell types and for each subtype. Several differentially expressed transcripts were identified across cell types (log2 fold change > 1 in absolute value, FDR < 0.05, Supplementary File S1). Interestingly, roughly half of these changes were in long non-coding transcripts (lncRNAs, Figure 2b). This represents a statistically significant enrichment relative to expectation (p < 10^-3^), with lncRNAs being at least three times more likely to be differentially expressed than other types of transcripts (Supplementary Table S5).

A similar pattern was observed at the subtype level. All analyzed cell subtypes exhibited transcripts with statistically significant differences between good learner and untrained conditions, ranging from 283 transcripts in EXC GLU-2 down to 6 in EXC IMN and ependymal cells. Consistent with the major cell-type analysis, lncRNAs were significantly overrepresented (FDR < 0.05) among differentially expressed transcripts in 9 subtypes out of the 13 considered for differential analysis (see Supplementary Table S6).

Numerous studies have identified molecular changes in the chick IMM associated with imprinting^7,10,12–22,30^. Our results recapitulate previous findings^40–43^ and associate them with specific cell types or clusters: the CHRM2 and CHRM3 genes, encoding muscarinic receptors, were differentially expressed in several glutamatergic subtypes, while the GABRG3 and GABBR2 genes, encoding GABA-A and GABA-B receptors, exhibited increased expression in good learners within EXC GLU-7, EXC GLU-3, and INH SST (Supplementary File S1).

We further examined whether the observed changes arise jointly across cell types or subtypes. Differential expression results showed a high correlation between log2 fold changes across all major cell types, with excitatory and inhibitory neurons showing the highest level of concordance (Spearman correlation coefficient ρ = 0.85, p < 0.01), while correlations with non-neuronal cells were slightly lower (ρ = 0.69 and 0.72 for glutamatergic and GABAergic cells, respectively, p < 0.01 in both cases). High correlation between log2 fold changes persisted at the subtype level (Figure 2c), suggesting that transcriptomic changes occur in a coordinated manner. In particular, glutamatergic and inhibitory subtypes form their own distinct clusters, with the exception of immature excitatory neurons (EXC IMN) which exhibited a partially divergent differential expression pattern (Figure 2c).

We then investigated whether transcriptomic changes are cluster-specific. Figure 2d reports the log2 fold-change for the twenty-five transcripts with the largest expression increases across all glutamatergic cells (|log2 FC| > 1, FDR < 0.05). When examined at the cluster level, they exhibit statistically significant changes (FDR < 0.05) only in specific subtypes, indicating that these subtypes are the primary drivers of the overexpression of these transcripts in the broader excitatory neuron population. Furthermore, Supplementary Table S7 lists the 257 transcripts that are differentially expressed in individual glutamatergic or inhibitory subtypes but not in their corresponding cell types, supporting the hypothesis that changes linked to imprinting may occur, or at least be more prevalent, in isolated clusters.

Taken together, our findings (a) highlight an overrepresentation of lncRNAs among differentially expressed transcripts, (b) provide insights into the specific cell types underlying the imprinting-associated molecular alterations reported in prior studies^17,40–43^, and (c) demonstrate that, although imprinting-related transcriptomic changes are broadly coordinated across cell types and subtypes, selected gene expression changes are specific to or predominantly driven by individual clusters.

### Imprinting is associated with biological processes related to neural plasticity

We next characterized biological processes (BPs) associated with imprinting through functional enrichment analysis across major cell types and neuronal subtypes (Supplementary File S2, Figure 3a). Across all major populations, cell–cell adhesion and cell junction organization were consistently enriched, indicating coordinated structural remodeling of the mesopallium circuitry during learning and corroborating previous findings^17^.

**Figure 3:**
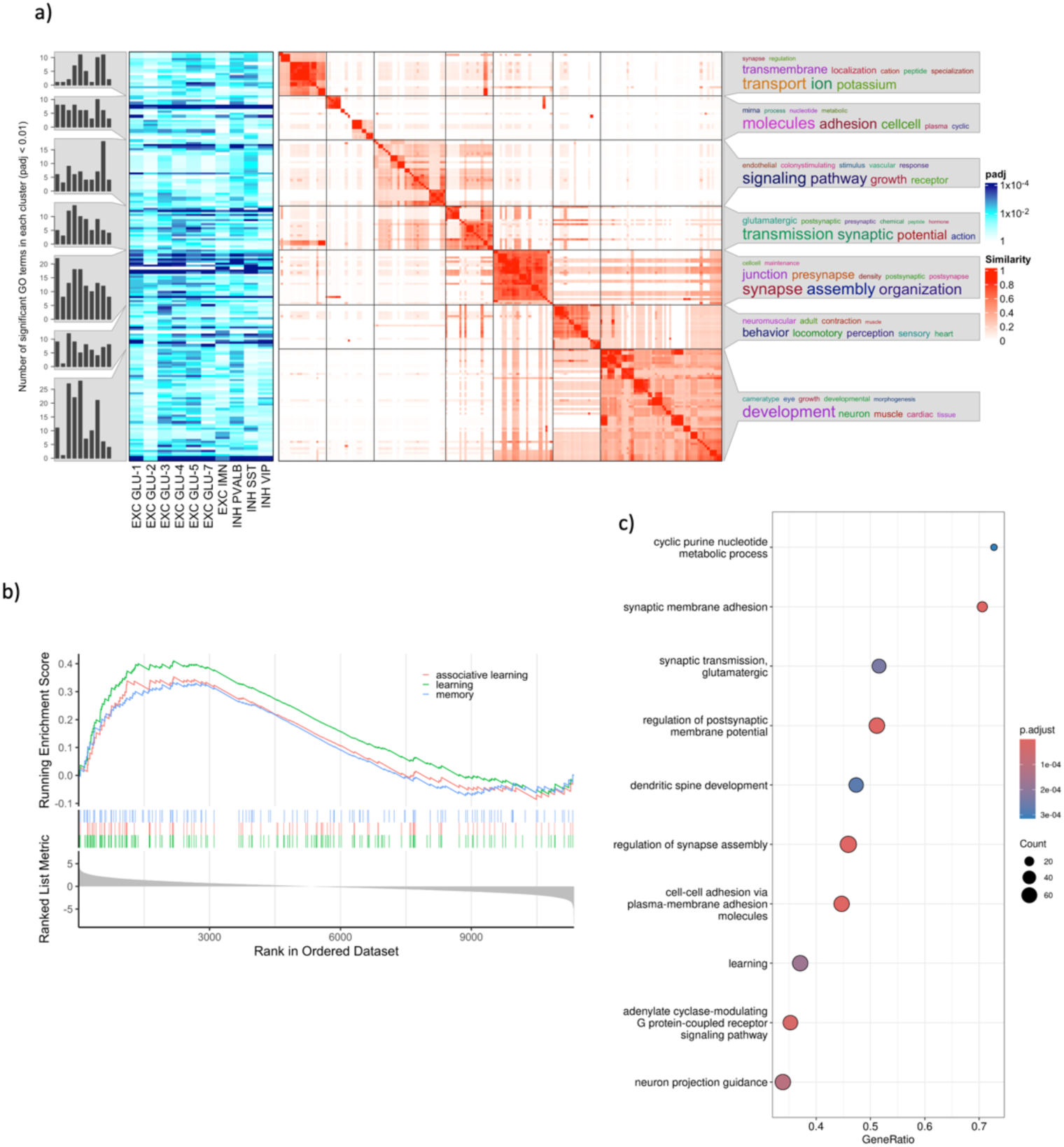
(a) Gene Ontology biological processes (BPs) significantly upregulated (FDR < 0.01) in good learner chicks, across neuronal subtypes, clustered and summarized. Central heatmap reports the BP-to-BP term similarity (see Methods), with BP clusters formed through the DynamicTreeCut algorithm. Left side heatmap reports the significance of each BP in each cell type (darker blue shades indicating higher significance). The word clouds on the right present the frequency of the most common terms within the BPs description. Bar plots on the left indicate the number of significantly upregulated BPs for each cell subtype and BP cluster. (b) Enrichment of relevant BPs in glutamatergic neurons: GO:0007612 learning, GO:0007613 memory, GO:0008306 associative learning. (c) Dot plot presenting the most upregulated BP in glutamatergic neurons; color indicate significance, size the number of upregulated genes within the BP.

Glutamatergic and GABAergic neurons exhibited convergent enrichment for synaptic membrane adhesion, postsynaptic signaling, and axon or neuron projection guidance, together with biological processes directly associated with higher cognitive functions, including GO:0007612 (“learning”), GO:0008306 (“associative memory”), and GO:0007613 (“memory”) (Figure 3b). Despite this convergence, the two neuronal classes displayed distinct functional specializations. Glutamatergic neurons were characterized by enrichment for excitatory synaptic transmission and cyclic nucleotide–dependent pathways (Figure 3c), consistent with potentiation and refinement of excitatory synapses. In contrast, GABAergic neurons showed pronounced enrichment for regulation of membrane potential, action potential, and ion transport processes, suggesting modulation of inhibitory neuronal excitability during learning.

Non-neuronal cell types displayed markedly different enrichment profiles. Astrocytes were primarily enriched for developmental and morphogenetic processes, consistent with a supportive and regulatory role in the learning-associated neural environment. Similarly, oligodendrocyte precursors were enriched for developmental patterning, cell fate commitment, and morphogenetic pathways, suggesting modulation of glial developmental programs accompanying imprinting.

We next examined imprinting-associated BPs at the neuronal subtype level. Among glutamatergic subtypes, EXC GLU-7 displayed a distinctive enrichment profile combining biological processes directly linked to learning and adult behavior with sensory-associated programs and regulation of neuronal excitability. The concurrent enrichment for sensory-linked processes, action potential generation, potassium transport, and adenylate cyclase–modulating GPCR/cyclic nucleotide signaling suggests that this deep-layer–like population integrates sensory-driven input with neuro-modulatory gating mechanisms relevant to imprinting-related circuit refinement.

In contrast, EXC GLU-2 was characterized by enrichment for dendritic spine development and vesicle fusion alongside prominent cyclic nucleotide signaling, indicating active structural and functional synaptic remodeling. Notably, GLU-2 uniquely showed strong enrichment for miRNA biogenesis and regulatory processes, pointing to an additional post-transcriptional regulatory layer that may contribute to the stabilization and persistence of learning-associated transcriptional states.

Across inhibitory neurons, the three analyzed subtypes shared a common backbone of synaptic adhesion and assembly processes but diverged along distinct functional axes. The PVALB population uniquely coupled contact remodeling with prominent adenylate cyclase–coupled GPCR/cyclic nucleotide signaling and regulation of pre- and postsynaptic membrane potential. In contrast, SST interneurons were most strongly defined by coordinated synapse assembly and intrinsic excitability programs, while VIP interneurons combined synaptic remodeling with the strongest axon or neuron projection guidance signatures and a clear neuro-modulatory GPCR/cAMP component.

Together, these enrichment analyses indicate that imprinting engages convergent but cell-type- and subtype-specific biological processes related to synaptic remodeling, excitability, and circuit reorganization.

### lncRNAs associated with imprinting map to cell-type-specific co-expression modules linked to synaptic and developmental processes

As significant changes in expression between good learners and untrained chicks appear to preferentially involve lncRNAs rather than protein-coding genes, we aimed to better understand the functions of these differentially expressed lncRNAs. To do so, we clustered all transcripts—including both protein-coding genes and lncRNAs – using a methodology designed for single-cell/nuclei RNA-seq (hdWGCNA^44^, see Methods). This analysis generated cell-type-specific transcript modules, encompassing 2,846 transcripts across 17 modules for glutamatergic neurons, 921 transcripts in 6 modules for GABAergic neurons, 3,476 transcripts in 6 modules for astrocytes, and 50 transcripts in a single module for OPC (see Supplementary Figure S16 and Supplementary File S3).

We then performed enrichment analysis on each module, considering only genes included in Gene Ontology BPs (Supplementary File S4). This approach allows assignment of biological functions to each module and, by extension, to the differentially expressed lncRNAs within them^45,46^. Differentially expressed lncRNAs associated with memory formation in glutamatergic chick neurons are distributed across multiple hdWGCNA modules, indicating functional heterogeneity. The largest contribution maps to a synapse-related module enriched for synapse assembly and regulation of postsynaptic membrane potential, suggesting lncRNA involvement in synaptic organization and excitability^47^. Additional lncRNAs associate with modules linked to dendritic morphogenesis, extracellular matrix proteoglycan metabolism, and developmental or regeneration-like neuronal programs, consistent with structural and circuit-level remodeling during learning^48^. A single DE lncRNA maps to a large mitochondrial/translation module, indicating a possible association with neuronal bioenergetic and proteostasis pathways. In astrocytes, three differentially expressed lncRNAs mapped to a single hdWGCNA module enriched for guidance- and adhesion-related processes. This module is characterized by genes involved in autonomic nervous system development and heterophilic cell–cell adhesion, including semaphorin–plexin–neuropilin signaling and synapse-associated adhesion molecules such as neurexins and neuroligins. These enrichments may point to astrocyte-mediated regulation of cell–cell interactions and circuit organization, consistent with an active role of astrocytes in synaptic remodeling during memory formation^49^.

Together, these findings indicate that both protein-coding genes and lncRNAs differentially expressed in good learners converge on cell-type-specific biological programs, prominently involving the same previously identified pathways on synaptic function, neural development, and circuit remodeling.

### Selected differential transcripts exhibit specific learning-related changes

Good learners show learning-induced molecular changes in the IMM, but they also differ from untrained chicks in other aspects, such as exposure to visual stimuli and levels of motor activity. Therefore, transcripts with prominent transcriptomic changes were further examined in experiments designed to isolate their specific role in memory, while controlling for potential confounding effects unrelated to learning. The following sections discuss the selection of the transcripts undergoing validation, as well as their respective results.

#### LUC7L, FOXP2, ROBO1, and RORA in learning and memory

Long-term memory formation involves specific and sustained transcriptional changes, mediated by transcription factors and other nuclear processes such as alternative splicing^50,51^. Based on statistical significance, fold change magnitude, and biological relevance, four differentially expressed protein-coding genes were selected for further analysis: two transcription factors (FOXP2 and RORA), a splicing-associated protein (LUC7L), and an axon guidance receptor (ROBO1) ^52,53^ . For all four genes, learning-related quantitative correlations were studied at the level of protein expression (the final product of gene expression).

The results of the immunoblotting experiments show that the levels of all four proteins were significantly (p < 0.05) correlated with preference score solely in the left IMM, except for FOXP2, for which a significant correlation was also observed in the left PPN (Figures 4 and 5, Supplementary Tables S8 – S11, Supplementary Figures S17 – S20). LUC7L, FOXP2, and RORA changes cannot be explained as a predisposition; they are rather attributable to learning during training. A similar finding holds for FOXP2 changes in the left PPN. Training had an additional effect on RORA expression, which was significantly depressed at preference score 50, indicating a suppressive effect of training unrelated to learning. ROBO1’s association with memory formation appears to be due to a predisposition, as the residual variance from the regression plot is significantly lower than the variance in untrained animals (P < 0.0005). The association with preference score is also weak, since there is no significant difference between the mean concentration of the protein in untrained chicks and the corresponding mean value in trained chicks with preference score 100 or 50.

**Figure 4.**
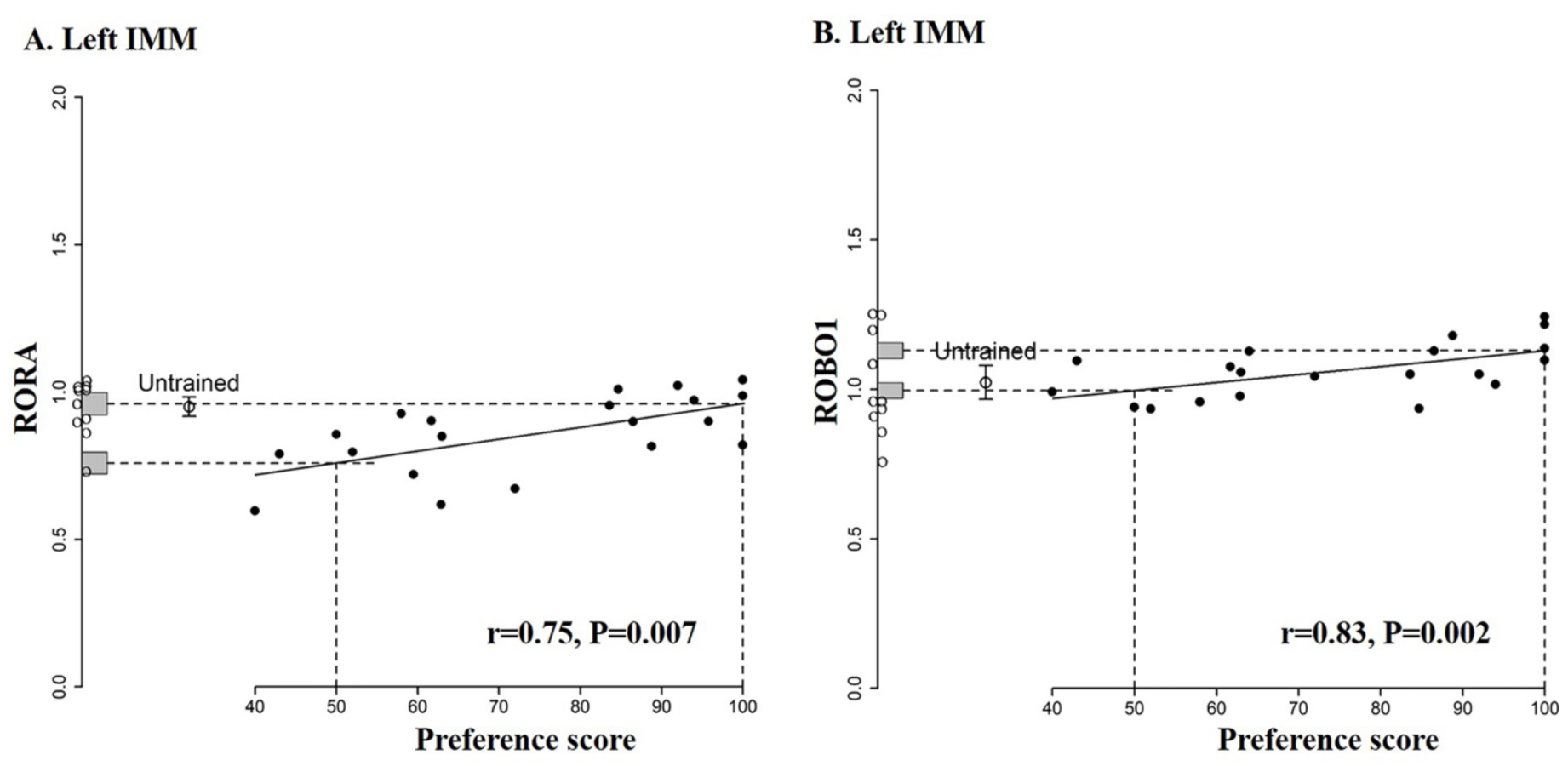
Standardized relative amount of RORA (A) and ROBO1 (B) proteins in the left IMM plotted against preference score. Filled circles, trained chicks. Open circle with error bars, mean level in untrained controls ± SEM; individual values for untrained chicks are shown as open circles along the y-axis. Vertical dashed lines, ‘no preference’ score 50 (approach to training and alternative stimuli equal/no learning) and ‘maximum preference’ score 100 (approach to training stimulus only/strong learning); horizontal dashed lines, y intercepts for ‘no preference’ and ‘maximum preference’. Gray bars on y-axis, ± SE of intercept. For both proteins correlations are significant

**Figure 5.**
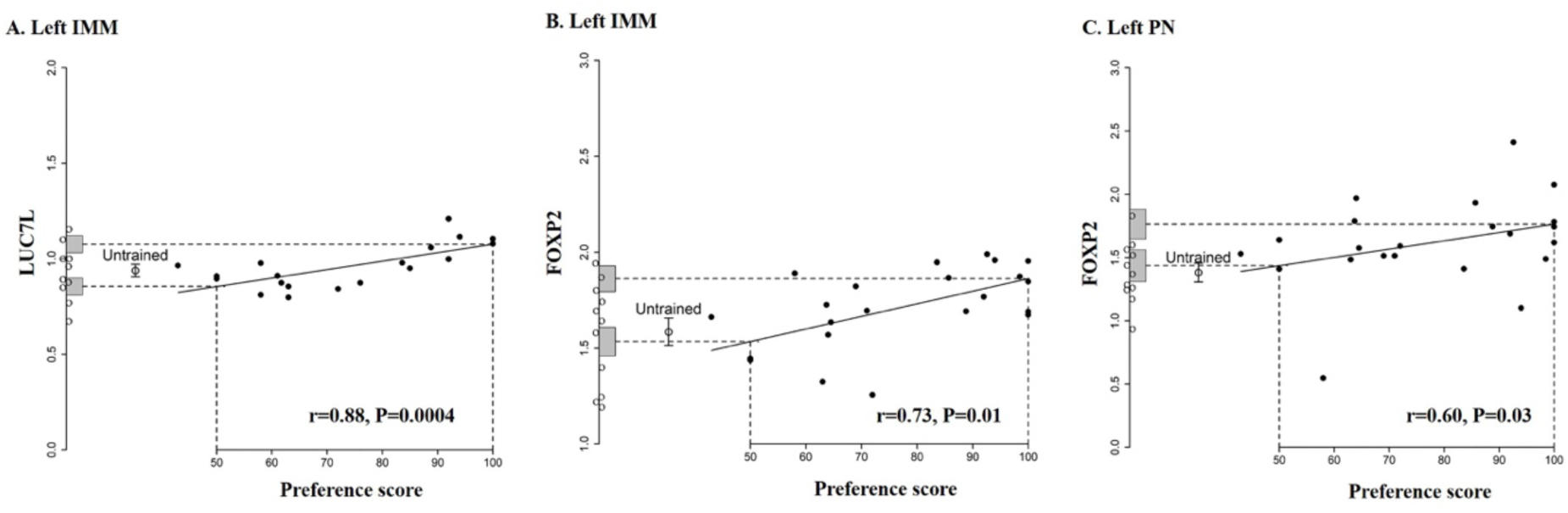
Standardized relative amount of LUC7L (A-left IMM) and FOXP2 (B-left IMM, C-left PPN) plotted against preference score. Filled circles, trained chicks. Open circle with error bars, mean level in untrained controls ± SEM; individual values for untrained chicks are shown as open circles along the y-axis. Vertical dashed lines, ‘no preference’ score 50 (approach to training and alternative stimuli equal/no learning) and ‘maximum preference’ score 100 (approach to training stimulus only/strong learning); horizontal dashed lines, y intercepts for ‘no preference’ and ‘maximum preference’. Gray bars on y-axis, ± SE of intercept. For all three cases correlations are significant.

#### Selected lncRNAs in learning and memory

Nearly half of the differentially expressed transcripts identified across all cell clusters corresponded to lncRNAs. On this basis, two lncRNAs were selected for further validation.

The first, ENSGALG00010007489, was the most strongly differentially expressed lncRNA in glutamatergic neurons. Notably, this effect was largely driven by a single subtype, EXC GLU-7, whose transcriptional profile resembles that of layer 6 (L6) neurons in the mammalian cortex, suggesting a highly cell-type–specific association. By contrast, the second selected lncRNA, ENSGALG00010026609 (hereafter referred to as lncRNA6609), displayed statistically significant differential expression across multiple major cell types, including excitatory neurons, inhibitory neurons, and oligodendrocyte precursor cells (Supplementary File S1).

We next examined the cellular context of ENSGALG00010007489 expression in greater detail. This lncRNA was predominantly expressed in cluster 2, one of the 5 clusters classified as EXC GLU-7 subtype (Supplementary Table S2), where it was detected in 22% of cells, compared with only 1.5% of cells across other glutamatergic clusters. Strikingly, 91% of cells in cluster 2 also expressed KCNMB2, compared with 21% in other glutamatergic neurons, representing a statistically significant enrichment (FDR < 0.001).

KCNMB2 encodes the β2 auxiliary subunit of large-conductance calcium- and voltage-activated potassium (BK) channels, which play key roles in regulating synaptic plasticity, neuronal excitability, and memory^54^. Based on this co-expression, we hereafter refer to ENSGALG00010007489 as GLUBK89.

In addition to BK2 channels (KCNMB2) and the GLUBK89 marker, the expression of several other genes distinguishes cluster 2 glutamatergic neurons from other glutamatergic populations (Supplementary File S5). Among the most significantly differentially expressed genes are MDFIC2, the transcription factors EST1 and ATOH8, and the contractile protein ACTN2. These gene expression differences contribute to alterations in multiple biological processes (Supplementary File S5). Notably, the most significantly affected biological process in cluster 2 neurons compared to other glutamatergic cells is regulation of endocytosis. This finding aligns with our previous work and that of others, which reported learning-related upregulation of proteins involved in endocytosis^16,55^.

We applied the same training protocol as for Western immunoblotting studies and standardized lncRNA (GLUBK89 or LNCRNA6609) expression levels were measured across all samples using qRT-PCR. Preference scores were recorded for all trained chicks (see Methods for details).

The results, presented in Figure 6A and Supplementary Table S12, show a significant correlation between GLUBK89 expression and preference score (Pearson’s r = 0.64, p = 0.03, Fig. 6A). The intercept at maximum preference is significantly higher than the mean expression in untrained chicks (p = 0.019). In contrast, the intercept at 50% preference (no learning) does not differ significantly from the untrained mean. Additionally, the residual variance from the regression is not significantly different from the variance in untrained chicks.

**Figure 6.**
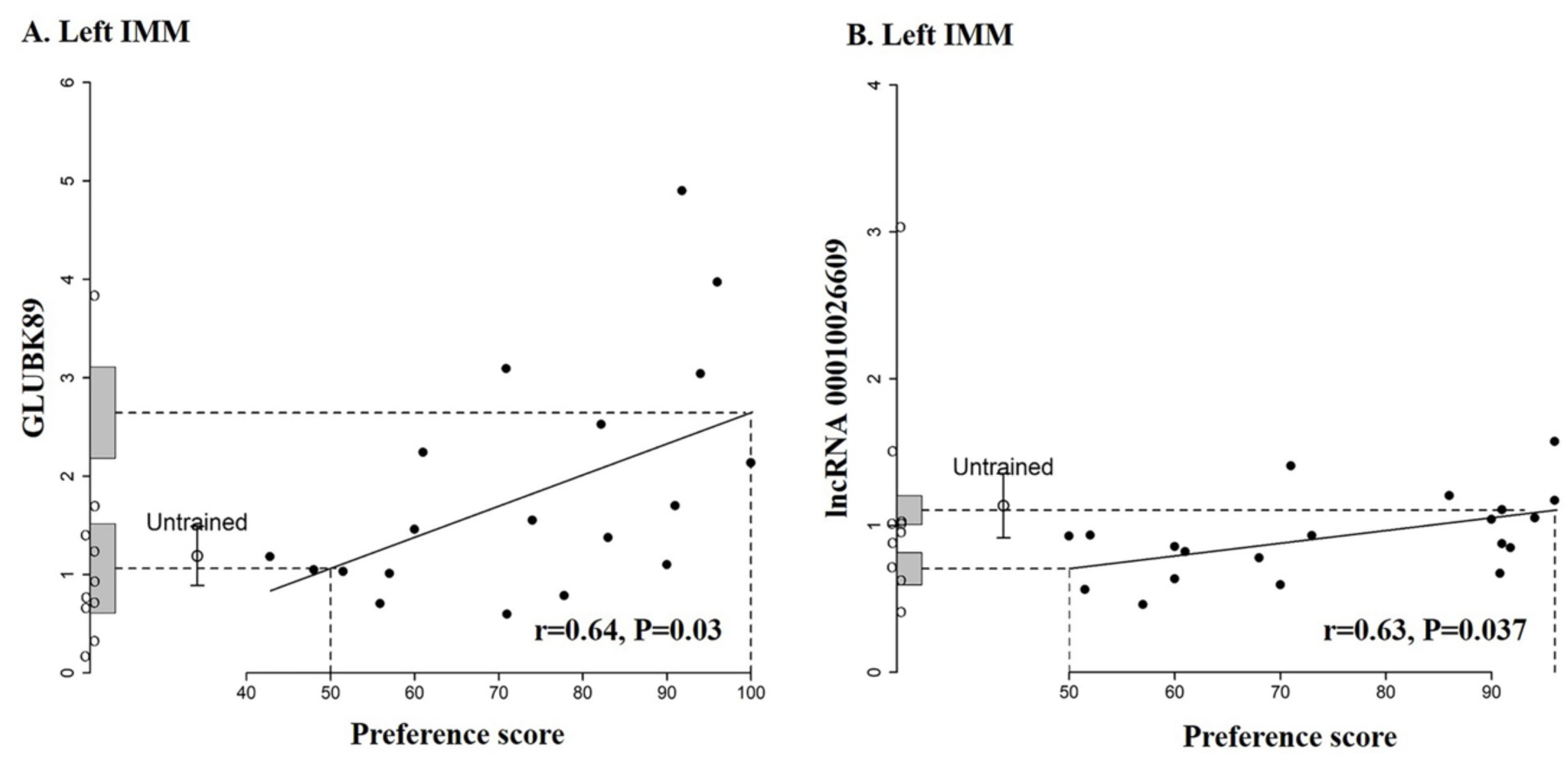
Standardized relative amount of GLUBK89 (A) and lncRNA 6609 (B) in the left IMM plotted against preference score. Filled circles, trained chicks. Open circle with error bars, mean level in untrained controls ± SEM; individual values for untrained chicks are shown as open circles along the y-axis. Vertical dashed lines, ‘no preference’ score 50 (approach to training and alternative stimuli equal/no learning) and ‘maximum preference’ score 100 (approach to training stimulus only/strong learning); horizontal dashed lines, y intercepts for ‘no preference’ and ‘maximum preference’. Gray bars on y-axis, ± SE of intercept. For both lncRNA the correlations are significant (P=0.03 and P=0.037 respectively).

These findings indicate that the observed positive correlation between GLUBK89 expression and preference score directly results from learning during training (see Methods). No significant correlations were found in other brain regions, suggesting a localized effect of learning in the left IMM (Supplementary Table S12 and Supplementary Figure S21).

The correlation between the standardized amount of LNCRNA6609 and the preference score is significant (*r* = 0.63, p = 0.037; Figure 6B, Supplementary Table S13). However, the differences between the mean untrained value and the intercepts for preference scores 50 and 100 were not significantly different. Together with the residual variance from the regression line being significantly lower than the variance of untrained chicks (p = 0.003), these results suggest that higher LNCRNA6609 expression may be associated with an inherent predisposition for learning rather than an effect of imprinting training (see Methods subchapter “Interpretation of a correlation between a molecular quantity and preference score”). No significant learning-related changes were evident in other brain regions (Supplementary Table S13 and Supplementary Figure S22).

#### GLUBK89 is a brain and nucleus specific transcript

In addition to their nuclear localization, lncRNAs can also reside in other cellular compartments, where they perform distinct functions. To determine the subcellular localization of GLUBK89 and LNCRNA6609, IMM tissue was fractionated into cytoplasmic and nuclear components, followed by RNA extraction and qRT-PCR analysis. GLUBK89 levels were found to be over 100-fold higher in the nuclear fraction, indicating predominant nuclear localization. LNCRNA6609 is present both in the nuclear and cytoplasmic fractions (Supplementary Figure S23).

Finally, the expression of both lncRNAs was analyzed in chick heart, muscle, liver, lung, and brain using quantitative RT-PCR. GLUBK89 expression was detected only in brain. In contrast, LNCRNA6609 was detected across all studied tissues, with the highest expression levels observed in the lung and liver (Supplementary Figure S24).

#### GLUBK89 is an avian-specific transcript, while LNCRNA6609 shows broad sequence conservation in mammals

To further characterize these lncRNAs, we examined their evolutionary conservation by identifying potential orthologs of GLUBK89 and lncRNA6609 in mammals.

Synteny analysis of protein-coding genes flanking the GLUBK89 locus on chicken chromosome 1 revealed a high degree of homology between chicken and turkey; however, chromosomal and regional conservation with the golden eagle, and especially with humans, was markedly reduced.

A similar analysis of the lncRNA6609 locus on chromosome 7 demonstrated strong conservation of gene content among chicken, turkey, and golden eagle, whereas conservation with the human genome was substantially lower (Supplementary Figure S25 and Supplementary File S6).

Further searches in the RNAcentral database revealed that GLUBK89 is conserved at the sequence level among avian species but lacks identifiable orthologs in mammals.

In contrast, lncRNA6609 shows sequence similarity with 89 regions in the human genome and 30 in the mouse genome. It shares more than 70% sequence identity with several human lncRNAs of unknown function, including HSALNT0096019 and SH3TC2-DT. Similarly, the mouse genome contains regions sharing over 65% identity with NONMMUT029999.2, NONMMUT009705.2, and NONMMUT032357.2, none of which encode proteins or have well-defined functions.

### FISH analysis confirms GLUBK89 localization in glutamatergic neurons and its association with memory

We further validated the cellular localization of GLUBK89 using triple fluorescence *in situ* hybridization (FISH). Specifically, we selected a good learner (preference score > 85%), a poor learner (preference score < 65%), and an untrained chick from each of four different batches, totaling 12 chicks. Six sections were collected from each brain. Five sections were triple-stained for GLUBK89, PPIB (a housekeeping gene), and SLC17A6 (a glutamatergic neuron marker). In the remaining section, PPIB was replaced with GAD2, a GABAergic neuron marker (see Supplementary Figure S26 for the experimental design). Regions corresponding to the left and right IMM and nidopallium were manually identified (see Methods for details). RNAscope images were processed using QuPath^56^ to obtain average fluorescence values for each channel within each detected nucleus (see Methods for additional details).

Figure 7 provides confocal FISH images of left IMM cells from a trained chick. Notably, the GLUBK89 fluorescent signal closely overlaps the glutamatergic marker SLC17A6 (panels A and C), while the same lncRNA is virtually absent from the GABAergic neurons marked with GAD2 (panels B and C).

**Figure 7.**
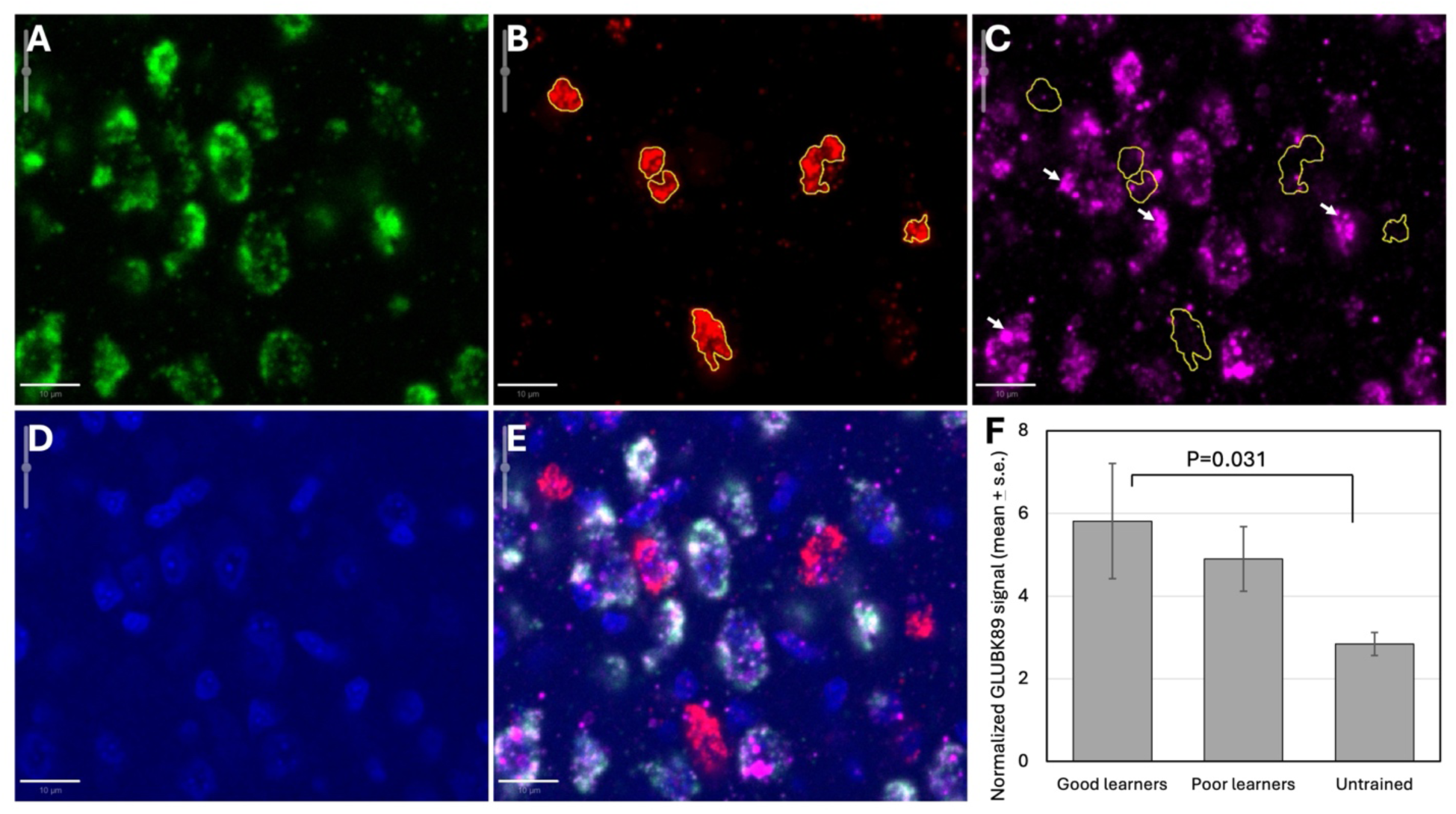
Representative confocal FISH imaging of the left IMM region in the brain of a trained chick. (A) Glutamatergic neurons expressing SLC17A6 are labeled with a green fluorescent marker. (B) GABAergic neurons expressing GAD2 are labeled with a red fluorescent marker, with their boundaries outlined in yellow. (C) GLUBK89 transcripts (purple) are primarily localized within the nuclei of glutamatergic neurons (indicated by arrows), rather than GABAergic neurons (outlined in yellow). (D) Nuclei are stained with DAPI. (E) Merged image. Bar=10 µm. (F) GLUBK89 expression, normalized by PPI, in the left IMM across groups.

To assess whether GLUBK89 expression in glutamatergic neurons is associated with memory of visual imprinting, we quantitatively analyzed sections stained with SLC17A6, GAD2, and GLUBK89 (Supplementary Figure S27) in different experimental groups of chicks. Nuclei of putative glutamatergic and GABAergic cells were selected as having above-threshold SLC17A6 (>1700) and GAD2 (>650) fluorescence levels, respectively. GLUBK89 fluorescence levels were consistently and significantly (two-tailed t-test p < 0.001) higher in glutamatergic than in GABAergic nuclei (Supplementary Figure S27).

In the IMM, there was a significant effect of experimental group (F2,6 = 6.42, P = 0.032); no significant effect of side and no significant interaction between experimental group and side were observed (P > 0.25). The mean value for good learners was significantly greater than that of untrained chicks (P = 0.031, adjusted by Tukey’s method for a family of three means). No other pairs of means differed significantly. The rank order of mean normalized values for lncRNA in SLC-positive nuclei was good learners > poor learners > untrained chicks (5.49, 4.76 and 2.68, respectively). This rank order is consistent with the result shown in Figure 6A, which was obtained with a larger sample and thus greater statistical precision.

The same analysis of variance model applied to data from the nidopallium showed no significant effects (Supplementary Figure S28).

Thus, the triple FISH experiments confirm that: (i) GLUBK89 is localized exclusively within the nuclei of glutamatergic neurons; (ii) its expression is non-uniform, and the low-frequency distribution indicates the presence of a distinct subpopulation of GLUBK89-expressing glutamatergic neurons; and (iii) elevated GLUBK89 expression within this subcluster is associated with learning. These observations are fully consistent with our snRNA-seq data.

## Discussion

The present study, for the first time, (i) characterizes the cellular and transcriptomic organization of the avian intermediate medial mesopallium (IMM), a brain region critical for learning and memory, at single-nucleus resolution, and reveals transcriptomic homologies with deep layers of the mammalian cortex; (ii) provides evidence for extensive involvement of long non-coding RNAs (lncRNAs) in long-term memory formation; (iii) identifies specific cell types and subcellular compartments underlying learning-related differential gene expression during imprinting; (iv) defines the biological processes associated with these differentially expressed genes; and (v) uncovers molecular signatures of visual imprinting in chicks at single-cell resolution. These signatures encompass both imprinting-specific mechanisms and molecular pathways shared with other forms of memory.

### IMM homology with mammalian brain structures and cell-type identification

The IMM is a key brain region involved in memory encoding during visual imprinting in chicks and is also essential for passive avoidance learning^57^. Visual imprinting involves recognition of a sensory stimulus, and analysis of its neural basis in chicks offers a powerful framework for studying recognition memory across species. Moreover, alterations in imprinting-related behavior in chicks have been proposed as a model for aspects of autism spectrum disorders^58^ and for long-term memory studies in general^3,6,7^.

Despite the widespread occurrence of imprinting-like processes in mammals and the fundamental importance of recognition memory, the correspondence between the avian IMM and mammalian brain regions has remained unclear. Early studies proposed that the IMM is homologous to the mammalian association cortex^8^. Subsequent work suggested similarities to neocortical layers 2 and 3 based on gene expression patterns^59,60^, while later analyses of transcription factor expression indicated that mesopallial excitatory neurons may resemble mammalian neocortical neurons located within layers 2, 3, 5, and 6^61^. In agreement with these earlier results, a recently published scRNA-seq atlas of the chicken brain draws parallels between mesopallium glutamatergic neurons and mammalian deep-layer cortical excitatory neurons using scRNA-seq, scATAC-seq, and regulatory data^31^. When the same atlas was used for classifying cells in our study, we found that 47.26% of excitatory neurons were classified with high confidence either as EXC GLU-7 or GLU-2, whose transcriptomic profiles are most similar to layer 6 (L6) and layers 4, 5 (L4/L5) of the mouse cortex, respectively.

Finally, a previous study reported that a substantial proportion of glutamatergic (VGLUT2+) neurons in the chick mesopallium co-express SATB2 and BCL11A, markers whose co-expression is characteristic of intra-telencephalic (IT) neurons in mammals^61^. Consistent with this, we identified co-expression of SATB2 and BCL11A in 20,714 IMM excitatory neurons (37.8%), constituting the majority of cells in several glutamatergic clusters (Supplementary Table S14; Supplementary Figure S29).

These findings further support the hypothesis that a substantial subset of IMM glutamatergic neurons shares molecular features with mammalian cortical excitatory neuron classes, including intra-telencephalic (IT) neurons.

### The IMM and learning-related molecular correlations

Visual imprinting is accompanied by behavioral and physiological side-effects, such as increased arousal and motor activity, which are not essential for learning or memory but may nonetheless be linked to biochemical changes. Such effects can confound the identification of molecular correlates that are specifically attributable to learning. An analytical framework has been developed to address this issue and to distinguish molecular changes arising directly from learning from those reflecting a pre-existing capacity or predisposition to learn^10^.

Lesion experiments^11^ have shown that, 26 h after training, a brain region outside the IMM (termed S′) can support retention of a preference for the training stimulus. Lesioning the right IMM approximately 3 h after training prevents the formation of S′, although the precise anatomical location of this region remains uncertain^3,6^. Importantly, despite disruption of S′ formation, chicks retain the acquired imprinting memory, indicating that the left IMM is required for memory retention at 24 h^62^.

Consistent with this interpretation, neurons in the intact IMM increase their selective responses to the familiar training stimulus (e.g., a red box or a rotating blue cylinder) over the 24 h period after training ^63–66^ A novel stimulus evoked only a baseline response in IMM neurons throughout the experiment. If chicks were allowed to sleep undisturbed 5-11 h after the start of training, at ∼20 h the chicks were strongly imprinted and neuronal responsiveness to the familiar imprinting stimulus was more than twice the pre-training level. If sleep was disturbed during the same critical time window, preference score showed no evidence of imprinting and IMM neuronal responsiveness collapsed to the pre-training level^65^. Manipulation of sleep over the day following imprinting training thus reveals a close association between learned responsiveness in IMM neurons and preference score. Learning-related increases in neural cell adhesion molecules (NCAMs) and clathrin heavy chain are observed at 24 h but not at 9.5 h after training^17,55^. Together with numerous other molecular changes reported at this time point in the left IMM^7^, these findings support the existence of a time-dependent cascade of biochemical events culminating in stable recognition memory. The functional roles of these lasting quantitative associations with memory, and recognition of the imprinting stimulus by IMM neurons, remain to be elucidated.

### Transcriptomic fingerprints of learning in the IMM

Single-nucleus transcriptomic analysis of the IMM in good learner and untrained chicks identified 319 differentially expressed genes in glutamatergic neurons of the left IMM (|log2 FC| > 1, FDR < 0.05). Remarkably, 180 of these transcripts were lncRNAs, highlighting their potential functional relevance in imprinting-related memory formation.

Several biological processes impacted by imprinting in the present study confirm results from our previous work. Cell adhesion processes, pathways of chemical synaptic transmission, dendritic spine organization, regulation of miRNA transcription, among others, are upregulated in the left IMM of good learner chicks. Specifically, our previous data indicate: (i) upregulation of NCAMs^17^; (ii) increased glutamate release^67^; (iii) elevated synaptic vesicle turnover (clathrin heavy chains, dynamin) ^16,55^; (iv) increased membrane bound MARCKS protein^68^ and (v) changes in mi-RNA spectrum^10^.

Although once considered transcriptional noise, lncRNAs are now recognized as key regulators of chromatin remodeling, transcriptional control, and post-transcriptional regulation ^69–71^. They function as molecular guides, scaffolds, and decoys, modulating chromatin architecture and gene expression through interactions with chromatin-modifying proteins and RNA-binding complexes, thereby contributing to cellular identity and genome organization.

Based on neuronal relevance and involvement in synaptic processes, we selected four protein-coding genes, namely LUC7L, FOXP2, ROBO1, and RORA, and two lncRNAs, GLUBK89 and lncRNA6609, for downstream validation.

Behavioral analyses combined with molecular measurements and established criteria for learning-relatedness revealed that GLUBK89, LUC7L, FOXP2, and RORA are upregulated as a consequence of learning at the protein level in the left IMM. In contrast, lncRNA6609 and ROBO1 were associated with a predisposition or capacity to learn rather than training-induced changes.

Detailed analysis of GLUBK89 using quantitative multi-probe *in-situ* hybridization (RNAscope), supported by RNA-seq data, demonstrated that its learning-induced upregulation is restricted to the nuclei of glutamatergic neurons. Although primarily expressed in a single cluster, GLUBK89 ranked highest in statistical significance in pseudo-bulk analyses across all glutamatergic neurons (Supplementary File S1). qRT-PCR analyses confirmed a learning-driven correlation exclusively in the left IMM, with no corresponding positive correlation in the right IMM or in either hemisphere of the PPN. GLUBK89 is further distinguished by its brain-specific, nuclear localization and apparent restriction to avian species. FISH analysis confirmed its localization to a small subset of glutamatergic neurons, supporting its role as an imprinting-specific regulator.

Given its co-expression with BK channels, known modulators of synaptic plasticity^72,73^, and its enrichment in a cluster characterized by KCNMB2 expression and endocytosis-related genes, we propose that GLUBK89 participates in coordinated gene regulation underlying learning and memory in this specific glutamatergic neuron population.

lncRNA6609, in contrast, is more broadly overexpressed across neurons and oligodendrocyte precursors. Unlike GLUBK89, lncRNA6609 is also localized in the cytoplasm and expressed in tissues beyond the brain. Its levels correlate with preference score exclusively in the left IMM, reflecting an inherent learning capacity rather than a training-induced effect, as inferred from variance partitioning criteria. Its cytoplasmic localization raises the possibility that lncRNA6609 functions through translation or as a scaffold for protein complexes, similar to previously described lncRNAs associated with four glycolytic payoff phase enzymes^74^.

LUC7-like proteins are components of U1 small nuclear ribonucleoproteins and regulate alternative splicing. Of the three vertebrate paralogs, LUC7L, LUC7L2, and LUC7L3^75,76^, only LUC7L showed learning-related differential expression. LUC7L expression was significantly higher in good learner chicks compared to the untrained group in almost all glutamatergic subtypes (EXC GLU-1, GLU-2, GLU-4, GLU-5, GLU-7), with a borderline significant increase observed in oligodendrocyte precursors. No significant changes were found for LUC7L2 or LUC7L3. Furthermore, we demonstrated that LUC7L protein levels positively correlate with learning strength specifically in the left IMM, while no such correlation was observed in other brain regions. This relationship is driven by the learning process itself rather than an inherent predisposition to learn.

LUC7L, LUC7L2, and LUC7L3 differentially regulate two major 5′ splice sites (5′SS) classes. The learning-related upregulation of specific splicing factor LUC7L suggests increased splice events during memory formation. This represents, to our knowledge, the first demonstration of memory-related regulation of LUC7L, consistent with emerging evidence implicating alternative splicing in synapse development and memory formation ^77^.

Retinoid-related orphan receptor-alpha (RORA) was upregulated in astrocytes and glutamatergic neurons of good learner chicks, with protein levels correlating positively with learning strength in the left IMM. This correlation was driven by the learning process rather than an inherent learning capacity.

In addition to this correlation, RORA expression at a preference score of 50 is significantly lower than in untrained chicks (Figure 4, Supplementary Table S10), indicating that the training procedure suppresses RORA expression in the absence of learning. This reduction could be attributed to various side-effects of training, such as locomotor activity, visual stimulation, or stress. In contrast, chicks that successfully learned during training exhibited increased RORA expression. Notably, at the highest preference scores (indicative of strong learning), the training-induced suppression of RORA was fully reversed. Given RORA’s versatility, it is perhaps unsurprising that its expression responds in opposite directions under different behavioral conditions, as illustrated in Figure 4.

RORA is a nuclear receptor family member that functions as a constitutive transcriptional activator^78^ and is predicted to regulate over 2,700 genes^79^. These include genes associated with autism spectrum disorders, neuronal differentiation, adhesion, synaptogenesis, synaptic transmission, cognition, memory, and spatial learning. Recent studies utilizing single-nucleus and spatial transcriptomics have identified neurons resilient to neurodegeneration in the human brain during Alzheimer’s disease progression. Some of these resilient neurons have been reported to be located in layer 4 of the association cortex and, early in the disease, exhibit upregulation of genes involved in synapse maintenance, synaptic plasticity, calcium homeostasis, and neuroprotection, including RORA and NCAM2^80^. Notably, the EXC GLU-2 subtype shares similarities with cortical layers 4/5 in mammals, and in our data these cells show upregulation associated with learning of both RORA (FDR = 0.007) and NCAM2 (nominal p-value = 0.012). To the best of our knowledge, this is the first demonstration of learning-related upregulation of RORA in a chick brain region known to be involved in memory.

Forkhead box protein P2 (FOXP2), a transcriptional repressor implicated in speech and cognitive functions^81^, was upregulated in GABAergic neurons, astrocytes, and oligodendrocytes of good learner chicks. FOXP2 may directly bind up to 400 promoters and regulate the transcription of at least 1800 genes in different tissues ^82^. Our snRNA-seq data reports the significant upregulation of FOXP2 in oligodendrocyte precursors (FDR = 0.02), as well as in astrocytes (FDR = 0.056). Further immunoblotting experiments revealed significant positive correlations between FOXP2 and the strength of learning and memory in the left IMM and left PPN. Both changes fulfill criteria of learning-relatedness^10^, and in both cases the up-regulation is attributable to training and rather than a predisposition to learn well. We speculate that learning-related changes in myelination may extend to the left PPN region. In all cases, the data obtained indicate long-term, memory-related changes of FOXP2-regulated biological processes.

This is the first time a change satisfying our criteria for learning-relatedness ^10^ (see Methods) has been detected in the PPN. The probability of all these criteria being satisfied by chance is low, raising the possibility that consequences of learning radiate from the IMM throughout the telencephalon in the hours following imprinting training. Indeed, regions outside the IMM can support memory for an imprinting stimulus under the control of the right IMM, in an area termed S’^8,11^, and several candidates for the location of S’ have been proposed ^83–85^. Whereas the critical role played by the IMM in the learning and memory of visual imprinting and passive avoidance memory is well established ^6,8^, there is no evidence that the PPN has any specific role in imprinting, as opposed to passively reflecting changes originating in the IMM. ROBO1 (roundabout guidance receptor 1) binds SLIT2 guidance molecules and is involved in axon outgrowth and orientation^86,87^. Importantly, in the avian brain differential regulation of SLIT–ROBO components has been linked to specialized forebrain circuits underlying learned vocal behavior, supporting a broader role for this pathway in experience-dependent circuit specialization^88,89^. Beyond development, ROBO1 expression has been detected in astrocytes and shown to increase in remodeling contexts such as forebrain ischemia, suggesting potential involvement in glia-mediated structural plasticity^90^. In our scRNA-seq data, the expression of ROBO1 is increased in good learner chicks in non-neural and GABAergic cells (FDR=0.066 and 0.087, respectively). ROBO1 protein amount strongly correlates with the strength of learning and memory only in the left IMM. Importantly, this correlation reflects a predisposition or capacity to learn rather than a result of training.

Our snRNA-seq and DEG analysis revealed a set of candidate transcripts, potentially involved in memory formation process. Of particular interest is the implication of lncRNAs. Some of the candidates have been analyzed in further detail, controlling for confounding influences by comparing untrained chicks with trained chicks that display behavioral evidence of particular levels of memory. This behavioral specificity was extended to the cellular level with GLUBK89, locating the learning-related changes in GLUBK89 in the nuclei of glutamatergic neurons in the IMM. Cellular analysis in conjunction with behavioral measurements is thus a powerful means of elucidating neural mechanisms of memory.

### Visual imprinting involves both specific and general molecular mechanisms of memory

Our findings demonstrate that avian imprinting engages both specialized, species-specific molecular pathways and conserved mechanisms shared across vertebrates. GLUBK89 represents an avian- and brain-specific lncRNA with a highly restricted cellular and subcellular expression pattern, suggesting a dedicated role in imprinting. In contrast, lncRNA6609 shows broad expression, mammalian homology, and correlation with learning capacity, indicating involvement in more general memory processes. This pattern suggests its potential involvement in broader forms of memory beyond imprinting and across species. These findings illuminate imprinting-specific molecular pathways while also contributing to our understanding of the general molecular architecture underlying memory.

### Limitations and further directions

This study lacks mechanistic insight into the specific roles of lncRNAs in memory formation. The poor sequence conservation of lncRNAs across species hampers the identification of mammalian orthologs and limits functional predictions. Moreover, current computational models for inferring lncRNA function are constrained by the scarcity of avian-specific data^91^. To elucidate the biological processes and molecular mechanisms involving these lncRNAs, genetically modified chicks with targeted gain- or loss-of-function perturbations would be highly informative. However, generating transgenic chicken models currently remains technically challenging. As an alternative, adenoviral-mediated delivery of lncRNA constructs could provide a viable approach. Complementary strategies, such as CRISPR-based genome editing or optogenetic manipulation of specific neuronal subpopulations may also help dissect the functional contributions of these lncRNAs to memory processes.

## Materials and Methods

### Chick training and apparatus

Fertile eggs (Cobb 500) were obtained from Sabudara poultry, Tbilisi, Georgia. In total 49 batches of eggs were incubated and hatched in darkness. Chicks were reared in isolation in darkness and at 22–28 h post-hatch trained for 1 h. During training each chick was exposed in a running wheel (1 revolution = 94 cm) to a training stimulus, a cuboidal red box rotating about a vertical axis. The box contained a light surrounded by a red filter (Lee Filters 106 Primary Red); the larger two sides of the box (18 × 18 cm) were translucent and vertical and the remaining sides (18 × 9 cm) were black. During training, the stimulus was rotating and illuminated for 50 seconds and then switched off for 10 seconds each minute. The maternal call (70 –75 dB) of a hen was played while the stimulus was on. This procedure strengthens imprinting to a visual stimulus^92^. As a chick attempted to approach the training stimulus, it rotated the running wheel and revolutions of the wheel were counted to provide a measure of approach activity (“training approach”). A preference test was performed without the maternal call 10 min after the 1 h training period ended. During the preference test, each chick in a running wheel was shown sequentially the training stimulus and an alternative stimulus that the chick had not previously seen, in the following order: training/alternative/alternative/training. Each exposure period during the test lasted 4 min and successive periods were separated by 4 min in darkness making 8 min exposure to each stimulus. The alternative stimulus was a right circular cylinder (height 18 cm and diameter 15 cm) with a translucent wall and vertical axis, rotating about this axis at 28 revolutions per minute. The cylinder contained a light surrounded by a blue filter (Lee filters HT 118 Brilliant Blue). See Horn (1989) for illustrations of the training and alternative stimuli^93^. A preference score was used to measure the strength of imprinting (i.e., learning), computed as (approach to training stimulus during test × 100/total approach during test). A preference score of ∼50 indicates poor learning whereas a score of ∼100 indicates strong learning.

Chicks from: (i) 3 batches were used for single-nuclei RNA experiments; (ii)10 batches for LUC7L measurements; (iii) 11 batches for FOXP2 studies; (iv) 10 batches for lncRNA ENSGAL000100007489 experiments; (v)10 batches for lncRNA ENSGALG00010026609; (vi) 10 batches for RORA measurements; (vii) 10 batches for ROBO1 and (viii) 4 batches for double in situ staining studies. Eight batches of chicks were shared for Luc7L and FOXP2 experiments, 5 batches were shared between the lncRNA experiments, 8 batches were shared between ROBO1 and RORa experiments. Thus, 49 batches of chicks were used for the reported data. For the single nuclei RNA-SEQ experiments, each batch consisted of one good learner and one untrained chick, except for one batch in which two untrained chicks were included. For all other experiments in each batch, there were up to three trained chicks and a control chick from the same hatch.

Chicks were decapitated 24 h after the end of training. For single-nuclei RNA experiments only left IMM was sampled, whereas for protein and RNA measurements IMM and PPN from the left and right hemispheres were collected. The locations of the IMM and PPN are described in Solomonia et al. (2013)^21^. Details of the IMM removal procedure are provided in Davies et al. (1985)^94^ and details of PPN removal in Solomonia et al. (1998)^17^. For double in-situ experiments whole forebrains were removed (see below). Samples were coded after collection and all further procedures were performed blind.

### Preparation of nuclear suspension

The nuclear fraction for RNA-SEQ was performed by a combination of methods described by Bakken et al., 2018 and Zhou et al., 2020. Frozen IMM from the left hemisphere was homogenized in a Nuclei PURE Lysis Buffer, (NUC-201, Sigma, formulated for solid tissues) supplemented with DTT, Triton X-100, protease inhibitors and 0.5% RNasin Plus RNase inhibitor (Promega). The procedure was carried out in a 1 ml Dounce homogenizer (Wheaton) using 10 strokes of the loose Dounce pestle followed by 10 strokes of the tight pestle to liberate nuclei. Homogenate was strained through a 30 μm cell strainer (Miltenyi Biotech) and centrifuged at 900 x g for 10 minutes to pellet nuclei. The supernatant was carefully removed. The pellet was resuspended in homogenisation buffer and strained through a 20 μm cell strainer. The filtered suspension was centrifuged at 900 x g for 10 minutes to pellet nuclei. Further separation of nuclei from myelin debris was carried our as described by Zhou et al, (2020). Nuclei pellets were resuspended in 500 uL Nuclei Wash buffer (1% BSA in PBS with 0.2 U/uL RNasin (Promega) and 900 uL 1.8 M Sucrose. This combined 1400 uL mixture was carefully layered on 500 uL 1.8 M sucrose and centrifuged at 13,000 x g for 45 min at 4 °C. The nuclear pellet obtained was strained through a 30 μm cell strainer, resuspended in Foetal bovine serum + DMSO (90% +10%) and stored for further experiments at -70 ^0^C.

### Single-nuclei RNA-seq library preparation and sequencing

Single cell transcriptome profiling was performed at SciLifeLab, Stockholm, Sweden. Frozen nuclei suspensions were thawed, washed in a small volume of PBS/BSA/RNase inhibitor at 500 g centrifugation and strained again. Libraries were prepared with a Chromium Controller and the Chromium Single Cell 3p RNA library v3 chemistry (10X Genomics, USA), according to the manufacturer’s instructions. The quality of the libraries was verified on a 2100 Bioanalyzer (Agilent Technologies, USA). Paired-end sequencing (read 1–28 bp, read 2–91 bp) was run on a NovaSeq 6000 (S2 flow cell) sequencer (Illumina, USA).

### Single-nuclei RNA-seq preprocessing

The 10x application CellRanger^95^ (version 7.1.0) was used for alignment and quantification of the raw sequencing files into count matrices. The option “--include-introns” was set to “true”, as recommended for single nuclei experiment, while the ENSEMBL GRCg7b.108 genome was used as reference. The resulting count matrices were then processed using several tools. DIEM (Debris Identification using Expectation Maximization)^96^ was used in order to identify and eliminate debris, meaning empty droplets most likely to contain ambient mRNA and no cell. DoubletFinder^97^ was then applied for removing droplets possibly containing more than one single cell. Finally, cells marked as debris by CellBender^98^ were also eliminated. The standard Seurat^99^ workflow was then used for removing cell with more than 5% of mitochondrial mRNA, as well as for normalizing and scaling the data. During scaling, we further normalize the data by regressing out of the expression value the amount of mitochondrial mRNA, the total number of reads and measured features. Integration across samples was performed using an “anchor” based approach^99^, using the 2000 most variable genes. The Leiden algorithm^100^ was finally employed on the integrated data for clustering the cells according to their transcriptomics profiles (the resolution value was set to 0.8).

### Cell type identification

The expression of known cellular markers was used in order to assign clusters to major cell types: ARPP21, SLC17A6, SV2B, SATB2 (glutamatergic neurons), SLC6A1, GAD1, GAD2, SLC32A1 (GABAergic neurons), OLIG2, SOX10 (oligodendrocytes), AQP4, GLI3, SLC30A12, and LRRIQ1 (astrocytes and ependymal). The Seurat R package was then sued for refining the cell type classification through label transfer^99^ from a reference atlas^31^.

Seurat’s label transfer is based on the concept of “anchors”, which are pairs of analogous cells between a well-annotated reference dataset and a query dataset; these anchors are found by identifying mutual nearest neighbors in a shared low-dimensional space, typically derived from PCA. Once anchors are selected, Seurat projects metadata (e.g., cell type labels) from the reference onto the query by weighting contributions from reference cells associated through the anchors.

The SAMap algorithm^38^ was then used to confirm the cell classification, using reference data from the mouse cortex and hippocampus^32^. In short, given two scRNA-seq datasets from different species, SAMap first identifies putative homologous genes through reciprocal BLAST searches. The homologous genes are then used for assessing the similarity, in terms of expression profiles, between cells across the two datasets. In this way, it is possible to transfer cell type annotations from one dataset to the other. In our analysis, we used SAMap default settings, performing BLAST searches on the proteome sequences. Only transferred annotations with SAMap score higher than 0.65 were considered.

### Single-nuclei RNA-seq statistical analysis

Possible imbalances in the proportion of cells originating from good learner and untrained chicks were assessed separately for major cell types and clusters, using a permutation approach^101^. Results were deemed significant for permutation-based FDR < 0.05 and ratio between the groups above 1.5.

Differential expression analysis between good learner and untrained chicks, corrected for sex, was performed on pseudo-bulk profiles. Given a specific cluster or cell type, pseudo-bulk profiles are derived by summing up for each gene the count values measured across the cells contained in that cluster / cell type. In this way a single expression profile is obtained for each sample (chick) out of several cells, simulating the same conditions obtained with bulk RNA-seq (hence the name). It has been found that pseudo bulk profiles significantly reduce false positive findings in differential expression analysis for single cell experiments^102^. We used the method based on the negative binomial distribution implemented in the DESeq2 R package^103^ to perform pseudo-bulk differential expression analysis separately for each cell type and cluster. Notably, DESeq2 forgo testing transcript that do not match specific criteria, for example if all expression values are equal to zero^103^. The differential analysis was forgone altogether if less than 50 cells were available for any chick for that specific cluster / cell type. Gene set enrichment analysis^104^ was performed separately for each cluster and cell type on the basis of the results of the differential expression analysis, using the clusterProfiler R package^105^. Gene Ontology (GO) Biological Processes (BP)^106^ were used as gene sets. The GO annotations for chicken have been known to be not as well curated as the annotations of other organisms^107^. Thus, we converted chicken genes to their mouse orthologs and used the GO mouse annotations for the enrichment analysis^108,109^. The simplifyEnrichment Bioconductor package^110^ was used for summarizing enrichment results across several analyses.

### Clustering snRNA-seq transcript with hdWGCNA

The hdWGCNA Bioconductor package allows to cluster transcripts from single cell and single nuclei RNA-seq experiments^44^. In short, hdWGCNA first groups individual cells within metacells^111^, which are supposed to represent cluster of cells sharing the same identical state. A single expression profile is then computed for each metacell, an operation that usually alleviates the drop-out problem common in single cell data^112^. Metacells were computed within each major cell type (Astrocytes, Oligodendrocytes, GABAergic and Glutamatergic neurons) and sample in our analysis. A matrix of transcript-to-transcript associations is then computed through a formula involving linear correlations elevated to a power coefficient. This formula is devised so that to emphasize strong correlations, and the power coefficient is identified automatically in a data-driven way. Finally, a dynamic tree cut algorithm^113^ is used to define strongly connected transcript modules. Transcripts that do not show strong connections are grouped in a default cluster, namely the “grey” module.

### BLAST analysis on lncRNA

We used BLAST (Basic Local Alignment Search Tool) to identify genes with significant sequence homologies to the Ensembl canonical transcripts of the lncRNA genes ENSGALG00010007489 and ENSGALG00010026609. BLAST analyses were conducted against chicken transcripts (cDNAs) and genomic sequences using default parameters. All hits were consolidated into a single list, and duplicates were removed.

### SDS Electrophoresis and Western Blotting

Brain tissue samples were rapidly homogenized in standard Tris-HC buffer containing phosphatase inhibitor and protease inhibitor cocktails. Sodium dodecylsulphate (SDS) solution was added up to a final concentration of 5% and the mixture was incubated at 95 °C for 3min. Protein concentrations were determined in quadruplicate using a micro bicinchoninic acid protein assay kit (Pierce). Aliquots containing 30 μg of protein in 30 μl of standard sample buffer were subjected to SDS gel electrophoresis and Western blotting^14^.

The proteins were transferred onto nitrocellulose membranes and stained with 0.1% w/v Ponceau S solution. The stained membranes were digitalized and analyzed for equal total protein loading with LabWorks 4.0 (UVP). The nitrocellulose membranes were then treated with PBS + 0.05% Tween 20 solution and a standard immunochemical staining procedure was carried out with peroxidase-labelled secondary antibodies and the Super-Signal West Pico Chemiluminescent substrate (Pierce, 34580). The following primary antibodies were used: anti-LUC7L (ABIN6745218, antibodies-online), anti-FOXP2 (ab1307, Abcam), anti-RORA (Invitrogen PA1-812), and anti-ROBO1 (Invitrogen PA5-29917). For each antibody, the detected protein band corresponded to the expected molecular weight: LUC7L 44 kDa, FOXP2 80 kDa, ROBO1 250 kDa, and RORA 60 kDa.

The optical densities of the target immunostained protein bands were measured with LabWorks 4.0 (UVP). Within each of the gel four internal protein standards (15, 30, 45, and 60 µg of total protein) were used for the calibration of autoradiographs. These protein standards were from IMM homogenate fraction isolated from control chick brains. We used the same proteins standards in all of the immunoblotting experiments performed. Optical densities of the immunostained internal standard bands increased proportionally with the amount of loaded protein: for all four proteins, plotting optical density against protein amount (15 – 60 μg) showed a virtually perfect fit to a straight line (Supplementary Figure S30).

The relative protein amount for each sample (e.g., LUC7L) was calculated by dividing the optical density of the sample band by the optical density corresponding to 30 µg of protein from the calibration curve. The values obtained by this normalization procedure are termed as the “relative amount” ^14^ of, e.g., FOXP2. There are differences between the batches resulting in variation in the protein amounts between them. To remove these variations the batch mean was subtracted from the relative amount of protein for each sample and this difference then added to the overall mean of all batches. This quantity is referred to as “standardized relative amount of protein”.

The densities of the target protein bands were not adjusted to the content of any cellular housekeeping proteins, such as glyceraldehyde 3-phosphate dehydrogenase or beta-actin, since it is not guaranteed that such reference proteins will be unaltered in our experiments^114–117^ Instead, we consistently verified equal loading of sample proteins by Ponceau S staining.

### RNA isolation and RT-PCR

Total RNA from chick brain samples were isolated with “Nucleospin RNA: DNA, RNA and Protein purification (740955 Macherey-Nagel). Complementary DNA (cDNA) was synthesized by High-Capacity cDNA Reverse Transcription Kit (4368814, Thermofisher Scientific). Relative ENSGAL000100007489 and ENSGALG00010026609 cDNA copy numbers were determined by real-time PCR using the Step One Real-Time PCR System (Applied Biosystems) with the SYBR Green detection method and was normalized to HPRT1^118^. The comparative CT (ddCT) method was used to determine the relative target quantity in samples^119^. The same fraction of IMM RNA isolated from the IMMs of untrained chicks was used in all experiments as a reference sample. Amplicons from randomly chosen samples were sequenced at BGI BIO Solutions Co., Ltd. (Hong-Kong, China). The primer sequences are provided in Table 1.

**Table 1:**
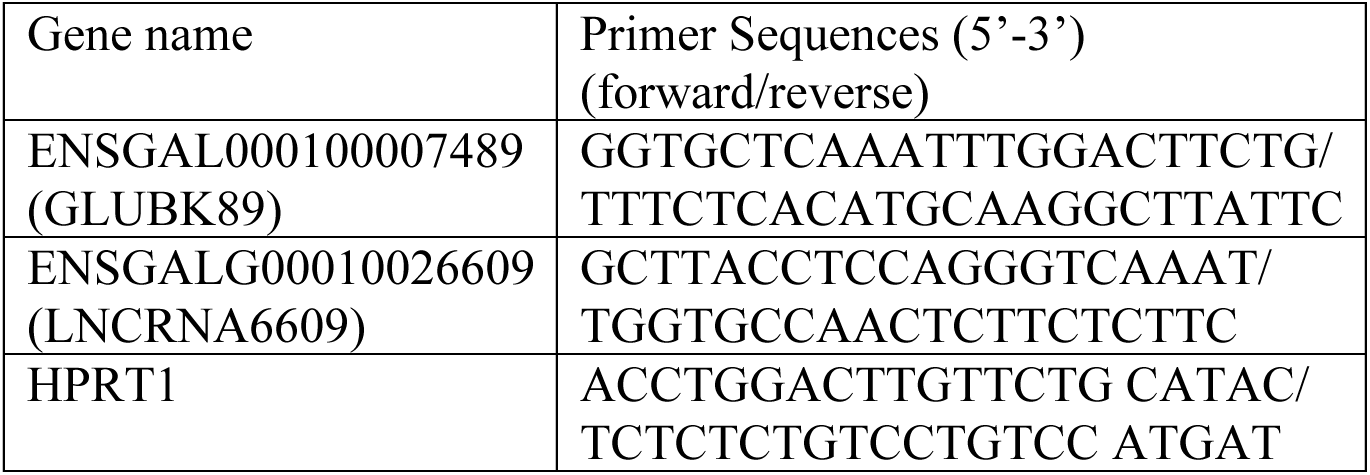
Primer sequences for RT-PCR.

### Subcellular localization of lncRNAs

Subcellular distribution of GLUBK89 and LNCRNA6609 lncRNAs was studied by a cytoplasmic and nuclear purification kit. (Cat. No. 21000 Norgen Biotek Corporation). The total RNA fraction was obtained separately from the nuclear and cytoplasmic fractions of the IMM and RT-PCR performed as described above.

### Interpretation of a correlation between a molecular quantity and preference score

A significant association between a molecular quantity (e.g., protein abundance, gene expression level, etc.) and preference score may indicate (i) that the change in the quantity is induced by the learning process, or that (ii) the quantity may facilitate / impede the learning process, i.e., it provides some type of predisposition. In order to distinguish between these possibilities, we use a design comprising two groups of chicks: a trained group, where both the molecular quantity under examination and the preference score are measured, and an untrained group, where only the molecular quantity can be measured. The molecular quantity is regressed over the preference score in the trained group using a linear model.

The y-intercept at preference score 50 is an estimate of the molecular quantity when no learning occurs in trained chicks. It thus measures the effect of procedures such as handling, locomotor activity, sensory stimulation etc. that occur during training but which are separable from learning (i.e. side-effects). If the intercept at preference score 50 is not significantly different from the mean value in untrained chicks, there is no evidence of side-effects.

In addition to a significant correlation with preference score, one would expect learning to give a y-intercept corresponding to the maximum preference score (100) that is significantly greater than the mean value for untrained chicks.

If the correlation with preference score is the *result* of learning (see (i) above), the residual variance about the regression line should be *statistically indistinguishable* from the variance in untrained chicks. This is because neither variance has a learning component but they are otherwise samples from the same population.

If the correlation is due to a predisposition (see (ii) above), training has no effect on the molecular quantity and the variance of the trained and untrained chicks should be statistically homogeneous. The correlation accounts for some of the variance in trained chicks and the variance about the regression line is correspondingly reduced: a significant *reduction* in residual variance about the regression line, relative to the variance in untrained chicks, is therefore evidence of a *predisposition*^10^.

### Fluorescent in-situ hybridization

The experiment was conducted in four batches. Each batch contained a good learner, a poor learner and an untrained chick from the same hatch of eggs, that is 12 samples in total. Hatching, rearing and training were performed as described for the other experiments. After killing, brain tissue was removed and fixed in 4% paraformaldehyde solution for 3 days. The brains were then transferred to a freshly prepared methacarn solution consisting of 60% (vol/vol) absolute methanol, 30% chloroform, and 10% glacial acetic acid^120^. Dehydration was performed through graded ethanol, followed by xylene, then paraffin wax for embedding. Samples were sectioned at 7 µm onto superfrost glass slides. For each brain, two coronal sections were selected corresponding to each of the coordinates A7.0, A7.6 and A8.0 of the chick brain atlas by Kuenzel and Masson^121^. Measurements of cellular fluorescence were made on each section within a rectangular sampling frame measuring 0.3 mm x 0.9 mm in the IMM and a square sampling frame 0.3 mm x 0.3 mm in the adjacent nidopallium (Supplementary Figure S31). Fluorescence was measured at each of the three anteroposterior levels for LOC121109133-O1, SLC and PPIB, and nuclei were stained with DAPI. One of the sections from each chick was stained for GAD instead of PPIB.

In any one batch, the sections stained for GAD were at the same anteroposterior level in all three experimental groups. There were three GAD sections from the posterior IMM in batch 1, three sections from the intermediate IMM in each of batches 2 and 3 and three sections from the anterior IMM in batch 4.

RNAscope was conducted to manufacturer specifications using probes Gg-GAD2-C1 (reference number 1300861-C1, target region 448-1421, 20 pairs), Gg-PPIB-C1 (reference number 453371, target region 67-876, 16 pairs), Gg-SLC17A6-C2 (reference number 1300871-C1, target region 591-1864, 20 pairs), Gg- LOC121109133-O1-C3 (reference number 1300881-C3, target region 91-656, 11 pairs), and Opal fluorophores 520, 570, and 650. Samples were scanned on a Zeiss Axioscan 7 at x20 and images were imported into QuPath^56^. Cell detection in QuPath was used, with the following parameters. Detection channel DAPI, requested pixel size 0.5 µm, background radius 8um, use opening by reconstruction yes, median filter radius 0um, sigma 1.5 µm, minimum area 10um², maximum area 400um², threshold 50, cell expansion 5 µm, include cell nucleus yes, smooth boundaries yes, make measurements yes, and visually validated. Subcellular detection in QuPath was then used, with the following parameters, detection threshold SLC 2000, PPIB 400, LNC 4000, smooth before detection yes, spilt by intensity yes, split by shape yes, expected spot size 1um², min spot size 0.1 µm², max spot size 4 µm², include clusters yes (still being optimized for all sections), and visually validated.

### Fluorescent in-situ hybridization data analysis

Quantitative measurements on fluorescent intensities provided by QuPath were analyzed using the R Statistical Software^122^. Two main analyses were performed:

- *Contrasting GLUBK89 expression values across experimental groups*. For this analysis, only sections stained for SLC17A6, PPIB and GLUBK89 were used (see Supplementary Figure 26 for a schema of the experimental design). Glutamatergic nuclei were identified by thresholding the SLC17A6 marker (> 1700), while a normalized expression value for GLUBK89 in each nucleus was computed by dividing for its fluorescent intensity for the fluorescent intensity associated to the housekeeping gene PPIB. These normalized expression values at the nucleus level were then averaged across sample, tissue (IMM and nidopallium), and side (left, right side of the brain). The resulting average values were modeled, independently in the IMM and nidopallium, according to the following linear mixed model^123^: norm_expression_values ∼ group * side, random = ∼ 1|batch/chick Which means that we model the normalized expression values depending on experimental groups (good learners, poor learners, untrained), side (left or right), and their interaction, as well as taking into account the variability provided by the nested random effect chick (sample) and batch (see Supplementary Figure S26 for batch definitions). The statistical significance of contrasts between experimental group levels within this model were investigated with the Tukey method^124,125^.
- *Contrasting GLUBK89 expression between GABAergic and glutamatergic cells*. Sections stained for SLC17A6, GAD2 and GLUBK89 were considered for this analysis. Glutamatergic and GABAergic nuclei were identified by thresholding the SLC17A6 (> 1700) and GAD2 (> 630) markers, respectively. The expression of GLUBK89 was contrasted between the two cell types using a t-test.

## Data and code availability

Data from the scRNA-seq experiments are available in Gene Expression Omnibus (https://www.ncbi.nlm.nih.gov/geo/query/acc.cgi?acc=GSE299793), while code for reproducing the results is available in GitHub (https://github.com/vlagani/CHARM-Vis).

## Funding

This work has been funded by (i) the European Union’s Horizon 2020 research and innovation program (project CHARM-Vis, project ID 867429), (ii) by the Shota Rustaveli National Science Foundation, project number NFR-22-8692, and (iii) by basic funding from Ilia State University and King Abdullah University of Science and Technology.

## Author contributions

V.L., R.S., Z.Kh., and D.C.G. conceived the project and designed the experiments. L.C., L.T., T.B., V.B., S.L., and A.J. performed the experiments. A.C.G.A., G.S., X.M.D.M., R.L., B.J.M., V.L. performed bioinformatic analysis and analyzed the data. V.L., R.S., Z.Kh., B.J.M., D.G.C., L.A.I., and J.T. interpreted the results and wrote the manuscript with input from all authors.

## Competing interests

The authors declare no competing interests.

## Notes

https://www.ncbi.nlm.nih.gov/geo/query/acc.cgi?acc=GSE299793

### Competing Interest Statement

The authors have declared no competing interest.

### Summary of Updates

Cell type identification revised, and subsequent analyses updated accordingly

