## Supplementary material for "Charting the single cell transcriptional landscape governing visual imprinting": Supplementary Material.pdf

### 1. Supplementary Figures

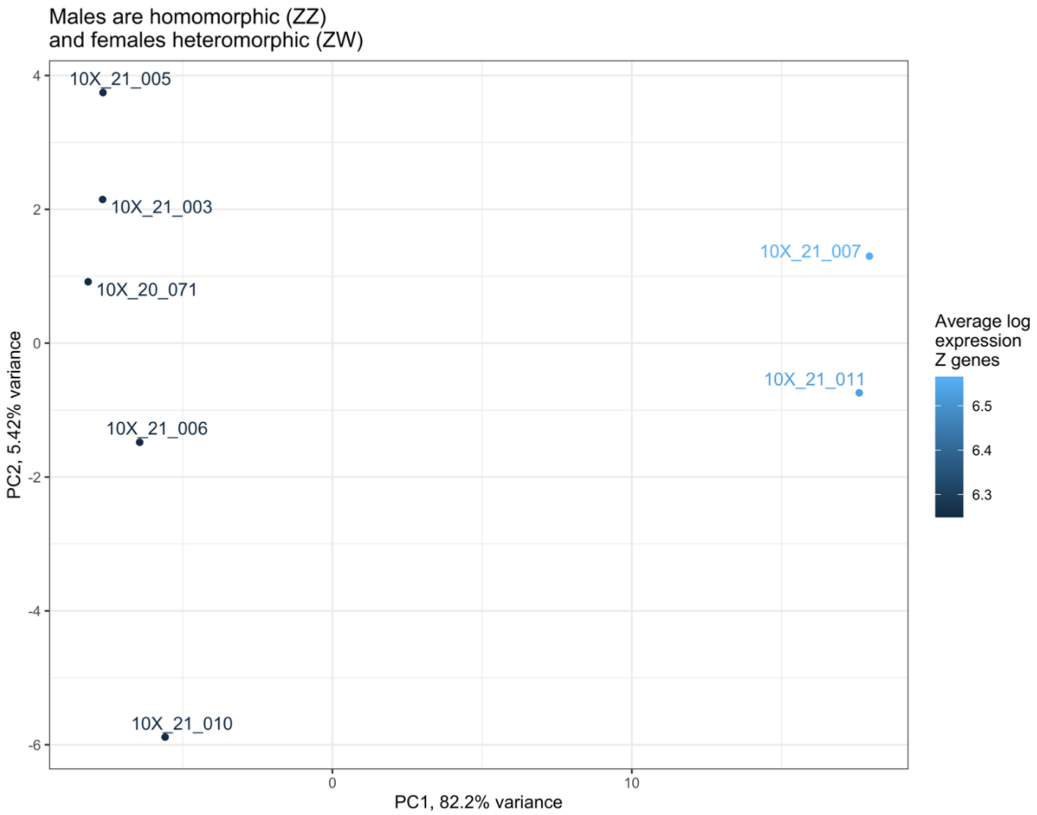

*Supplementary Figure S1: PCA analysis performed on the normalized, pseudo bulk expression profiles of the seven snRNA-seq samples, using only the gene in the sex chromosomes. The first component clearly separate male and female chicks, with the male chicks (right) showing a higher average expression of the genes in the Z chromosome.*

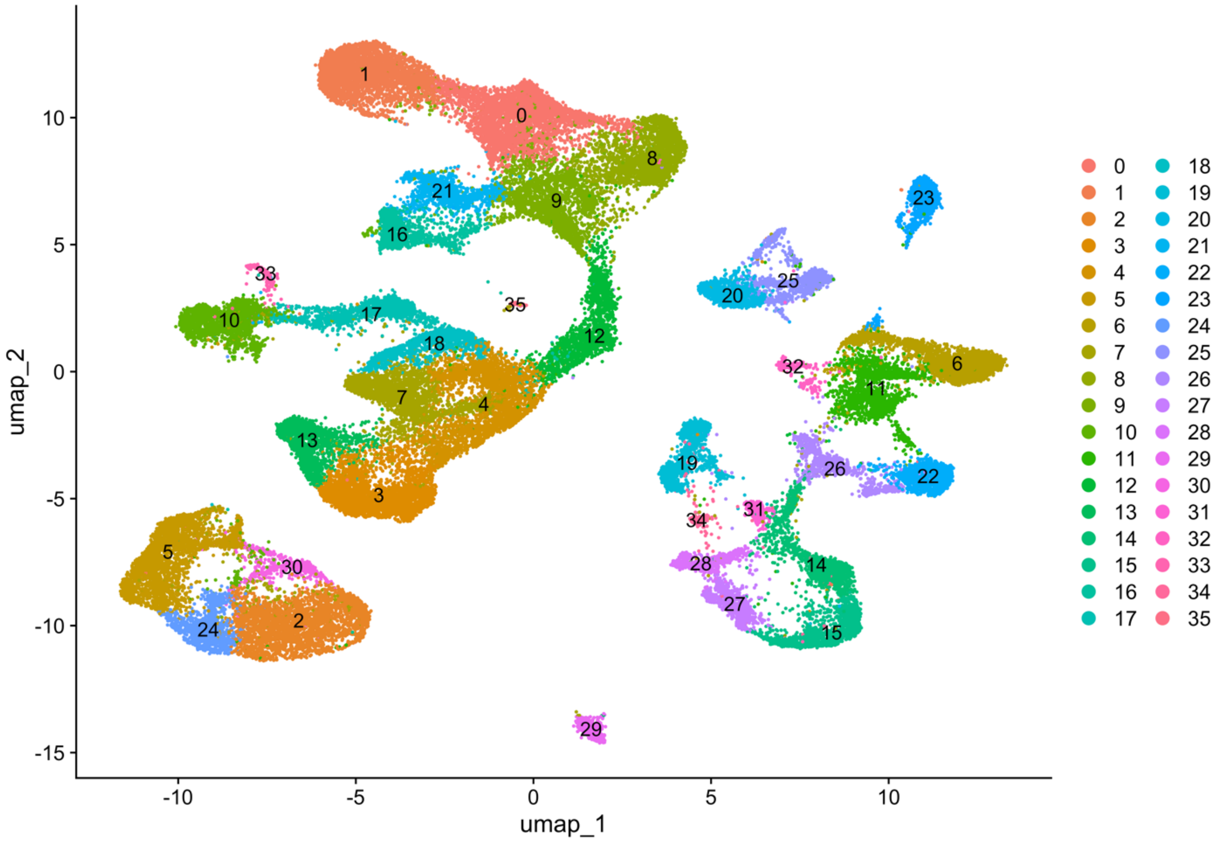

*Supplementary Figure S2: UMAP presenting the subdivision in clusters of all the 54763 nuclei included in the analysis across all samples. Different clusters are identified by color and label.*

### 1.1 Markers for the identification of GABAergic subpopulations

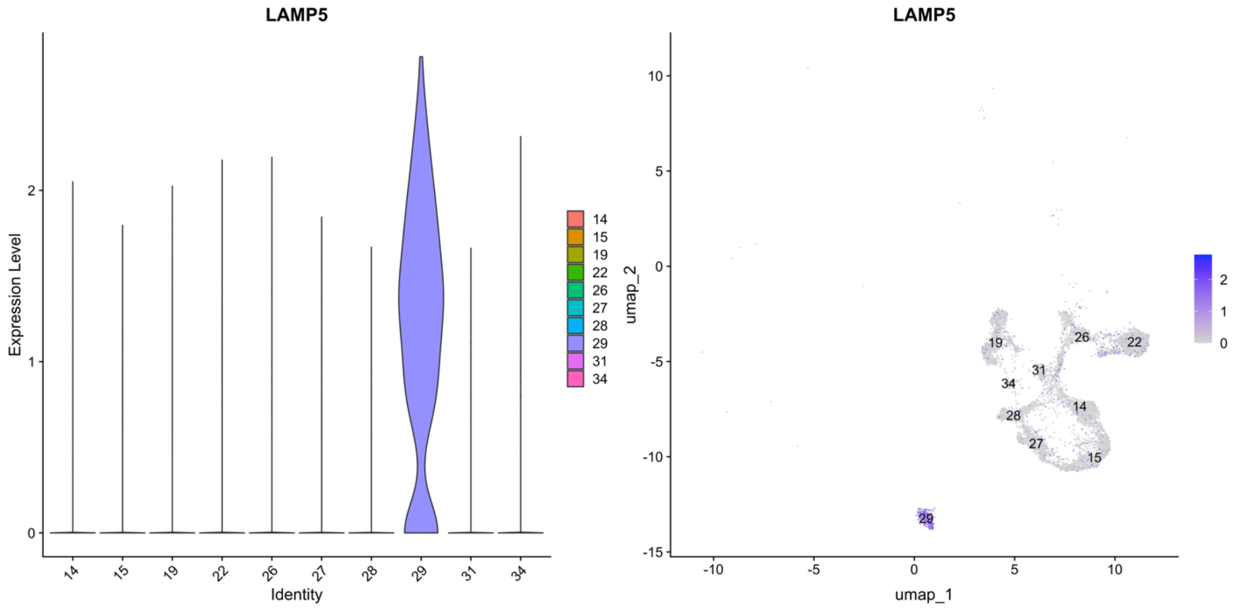

*Supplementary Figure 3: violin and UMAP plots for the gene LAMP5, marker for the LAMP5 GABAergic subtype.*

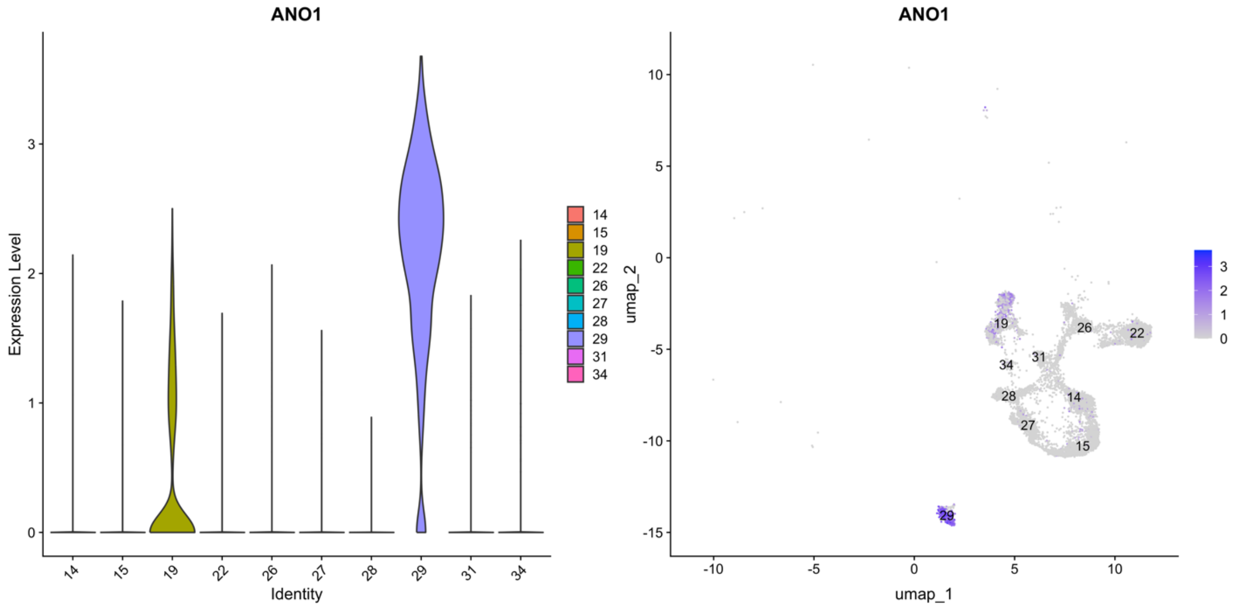

*Supplementary Figure 4: violin and UMAP plots for the gene ANO1, marker for the LAMP5 GABAergic subtype.*

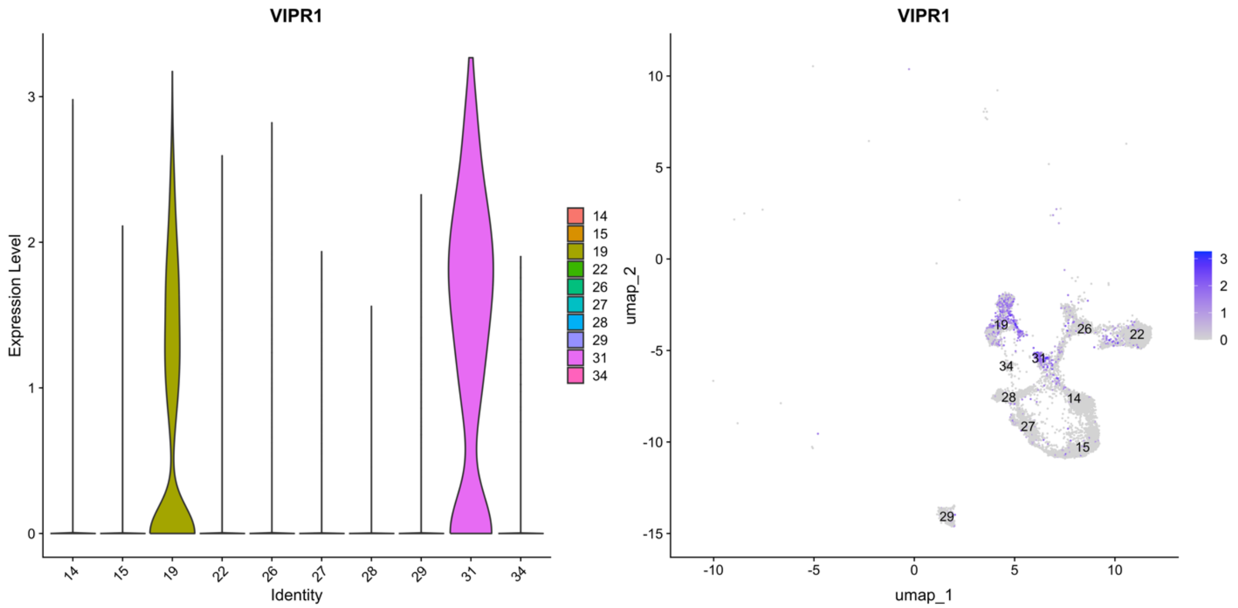

*Supplementary Figure 5: violin and UMAP plots for the gene VIPR1, marker for the VIP GABAergic subtype*

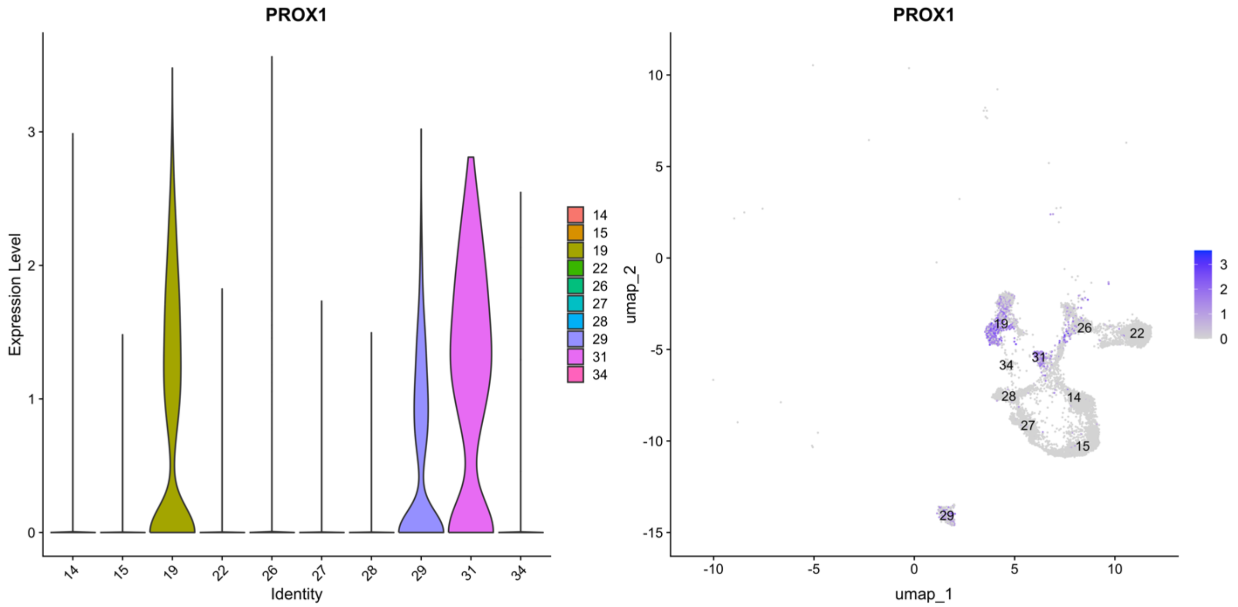

*Supplementary Figure 6: violin and UMAP plots for the gene PROX1, marker for the VIP GABAergic subtype*

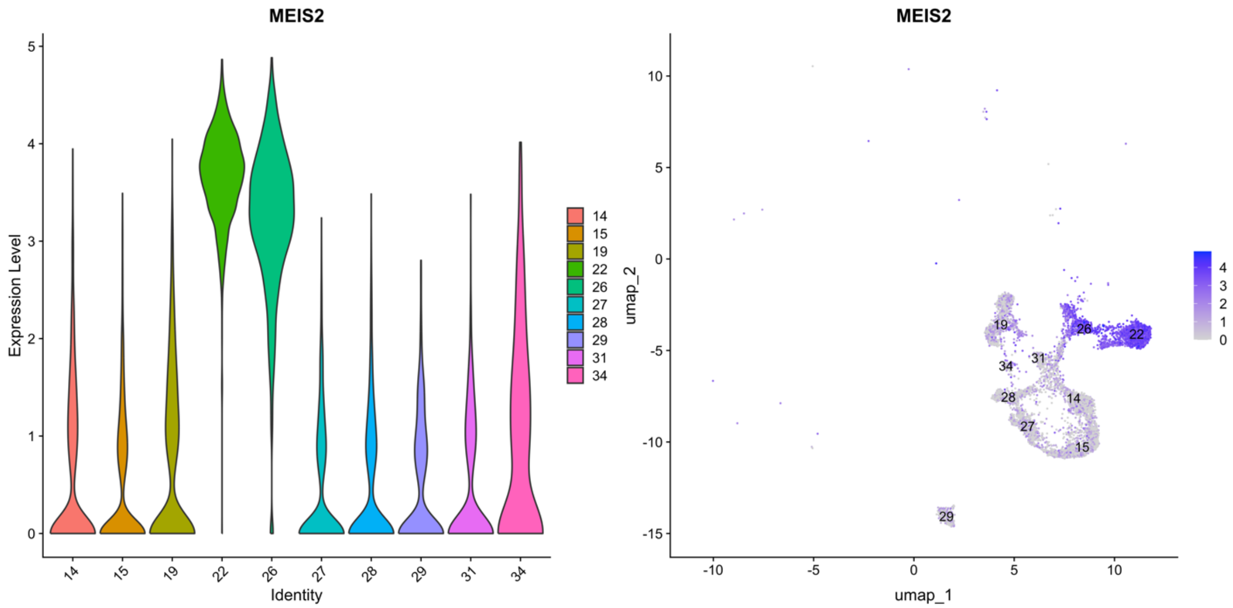

*Supplementary Figure 7: violin and UMAP plots for the gene MEIS2, marker for the MEIS-COL12A1 GABAergic subtype*

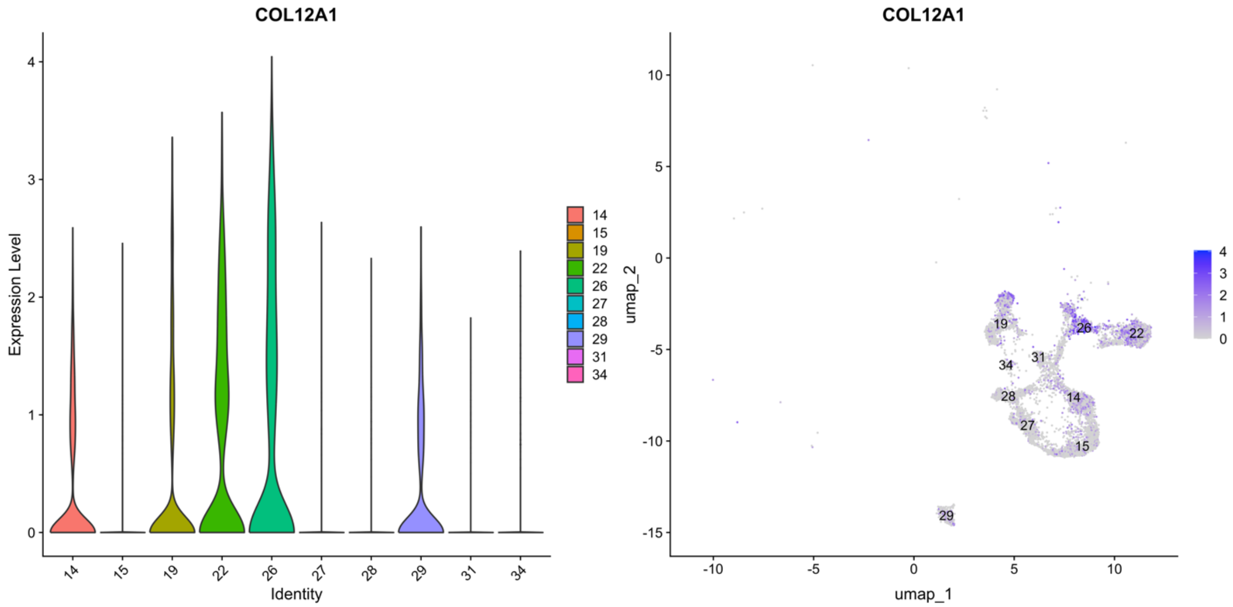

*Supplementary Figure 8: violin and UMAP plots for the gene COL12A1, marker for the MEIS-COL12A1 GABAergic subtype*

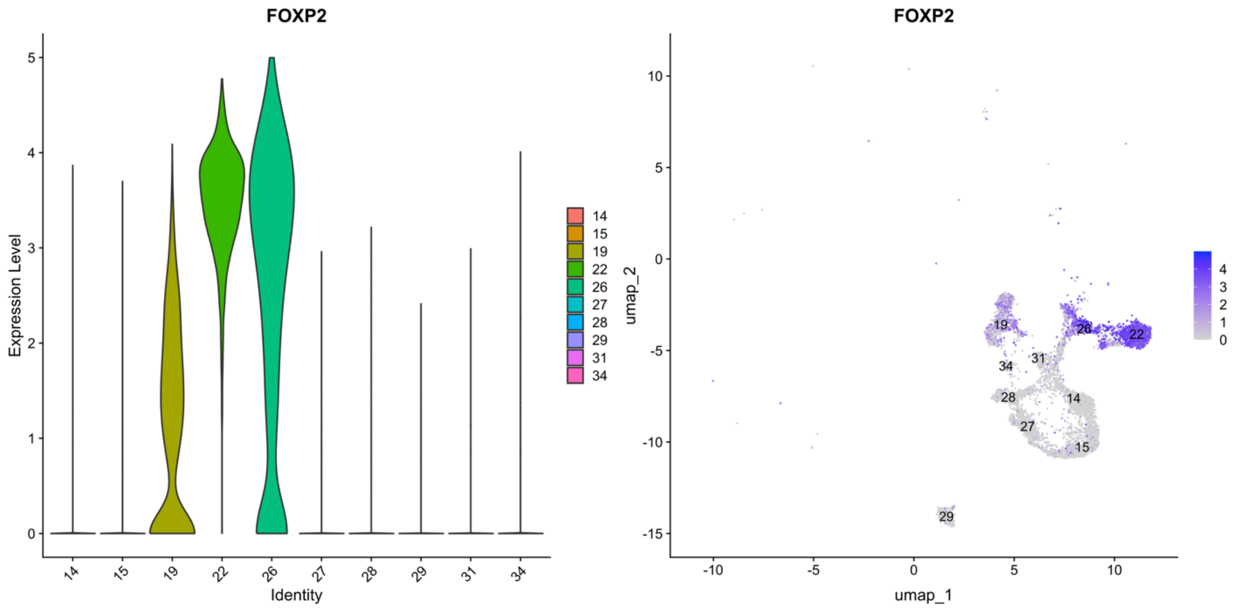

*Supplementary Figure 9: violin and UMAP plots for the gene FOXP2, marker for the striatal-like inhibitory neurons (STR) GABAergic subtype*

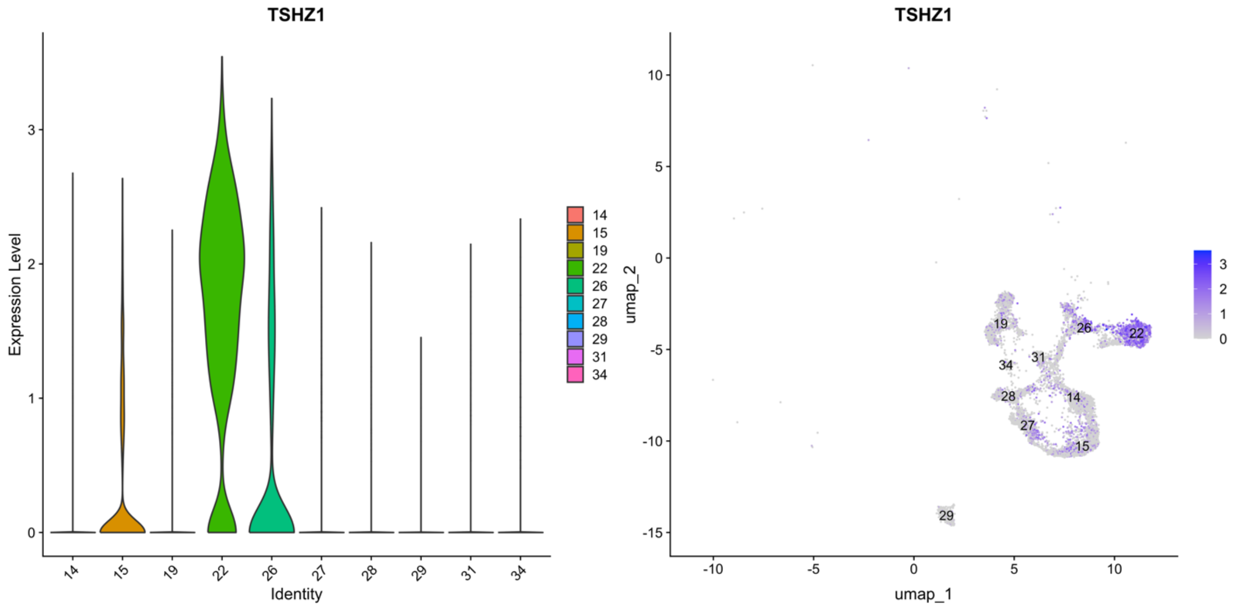

*Supplementary Figure 10: violin and UMAP plots for the gene TSHZ1, marker for the striatal-like inhibitory neurons (STR) GABAergic subtype*

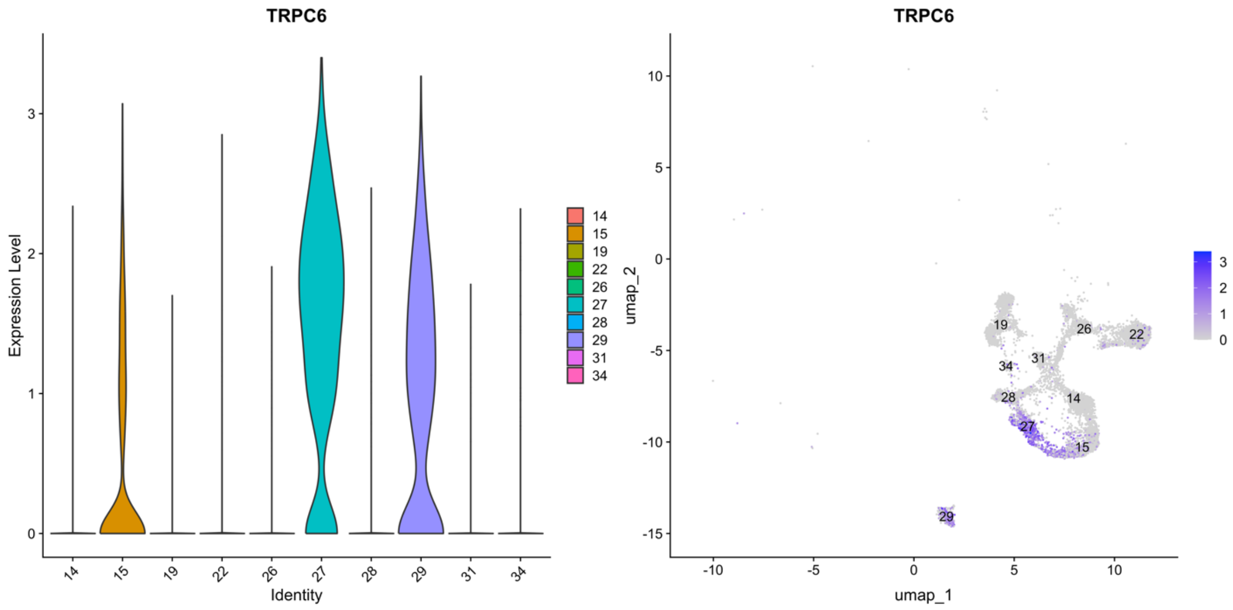

*Supplementary Figure 11: violin and UMAP plots for the gene TRPC6, marker for the SST GABAergic subtype*

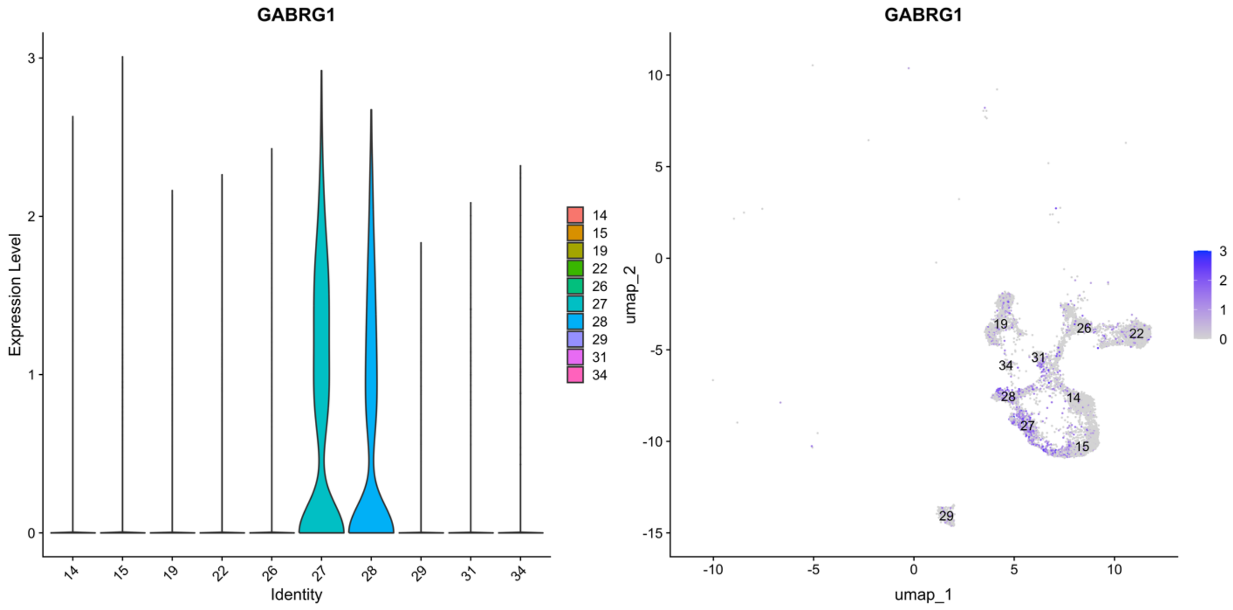

*Supplementary Figure 12: violin and UMAP plots for the gene GABRG1, marker for the SST GABAergic subtype*

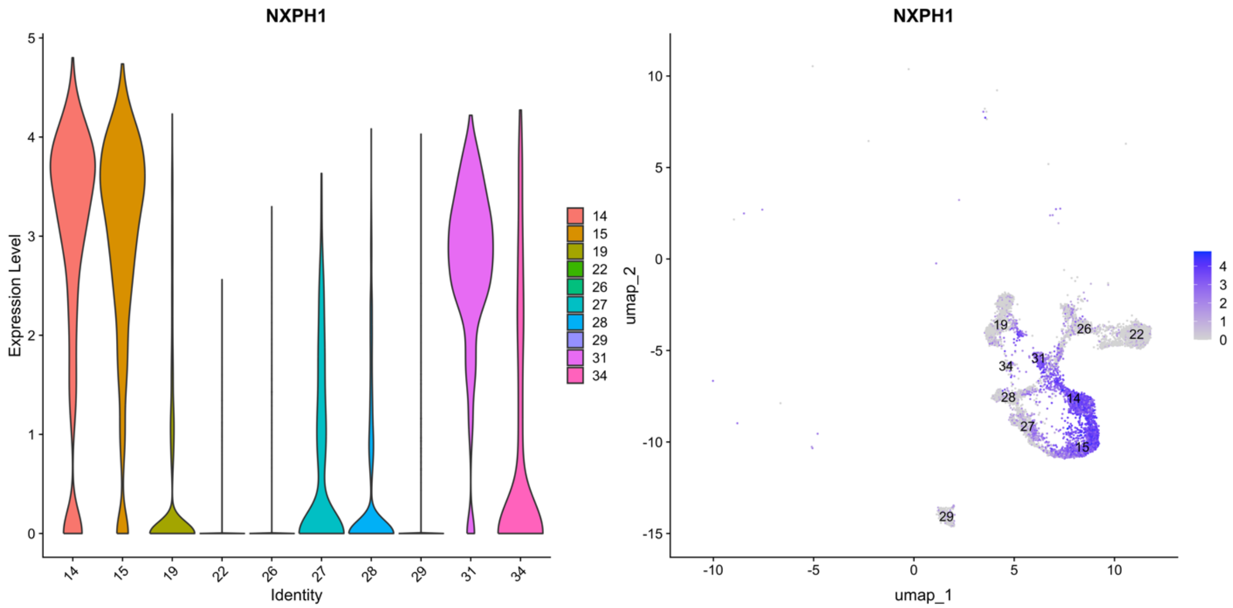

*Supplementary Figure 13: violin and UMAP plots for the gene NXPH1, marker for the PVALB GABAergic subtype*

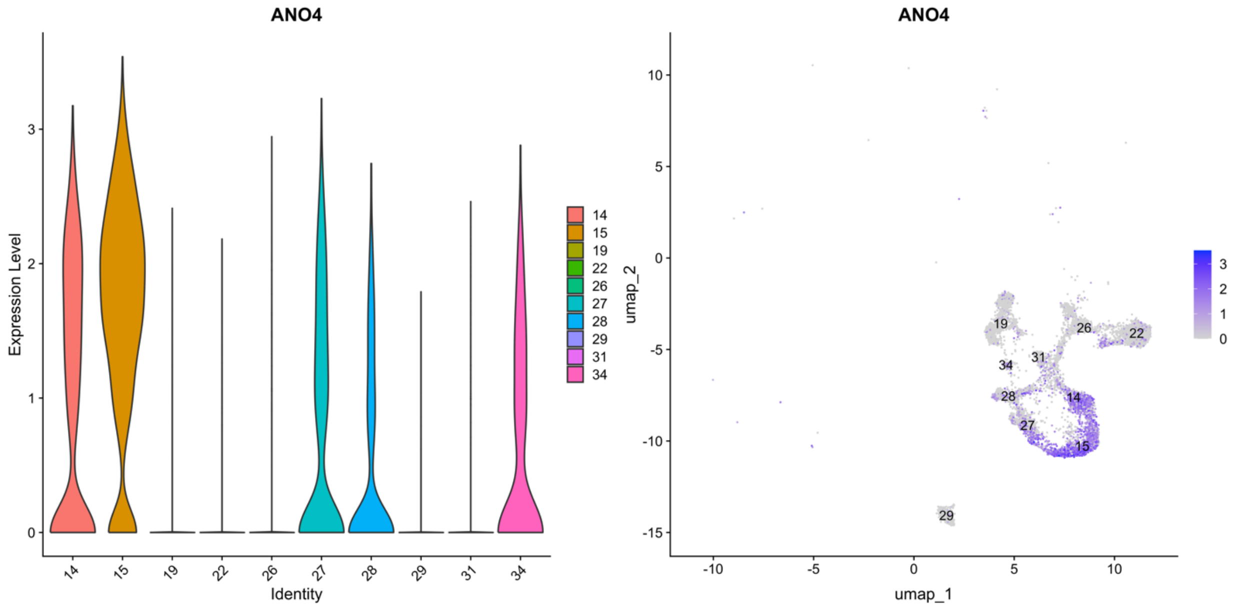

*Supplementary Figure 14: violin and UMAP plots for the gene ANO4, marker for the PVALB GABAergic subtype*

### 1.3 Figures related to transcriptomics changes between good learner and untrained

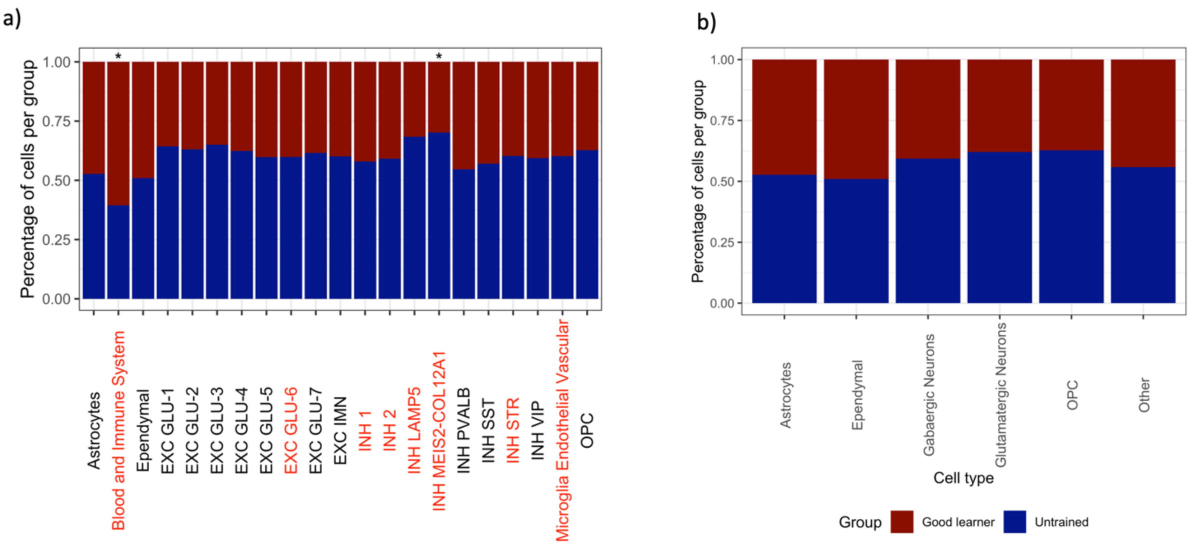

*Supplementary Figure 15: Percentage of cells from good learner (dark red) and untrained (dark blue) samples. Panel a): percentages across cell subtypes. Asterisks above the bar indicate clusters where the disproportion between the two groups is statistically significant (false discovery rate < 0.05, ratio > 1.5). Red numbers along the x-axis indicate the subtypes that have been excluded from differential analysis, as the number of cells in the least represented sample is less than 50. Panel b): percentages across major cell types. No major cell type had a statistically significant disproportion across the two experimental group, and for each cell type the least represented sample includes at least 50 cells.*

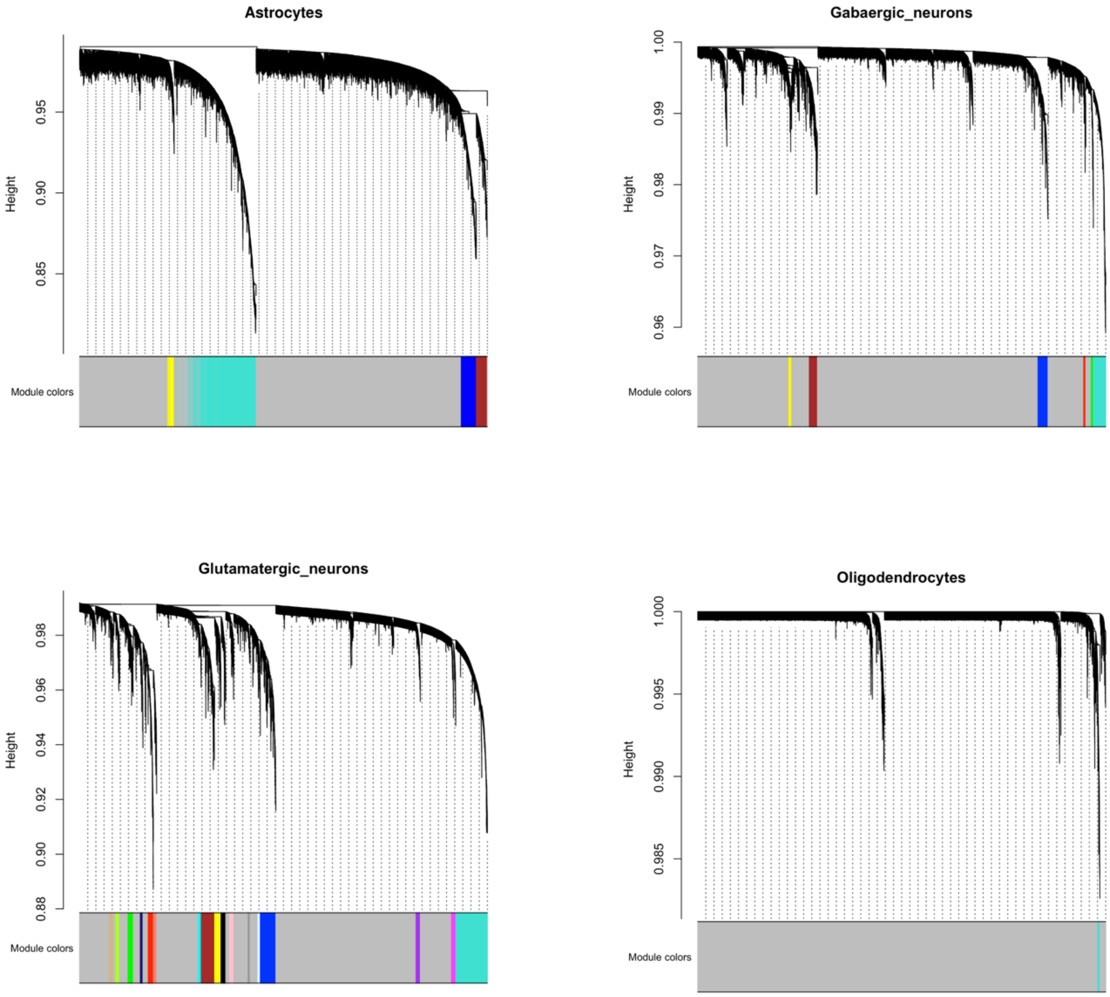

*Supplementary Figure 16: dendrograms representing the transcript modules produced by the hdWGCNA algorithm. Each panel corresponds to a different cell type. Transcripts lay on the x-axis, while the y-axis reports the similarity between transcripts (the lower the branch of the dendrogram, the strong the association). Modules identified by the dynamic tree cut algorithms are represented with colors, while transcripts not included in any clusters are in grey.*

### 1.4 Figures related to experimental validation

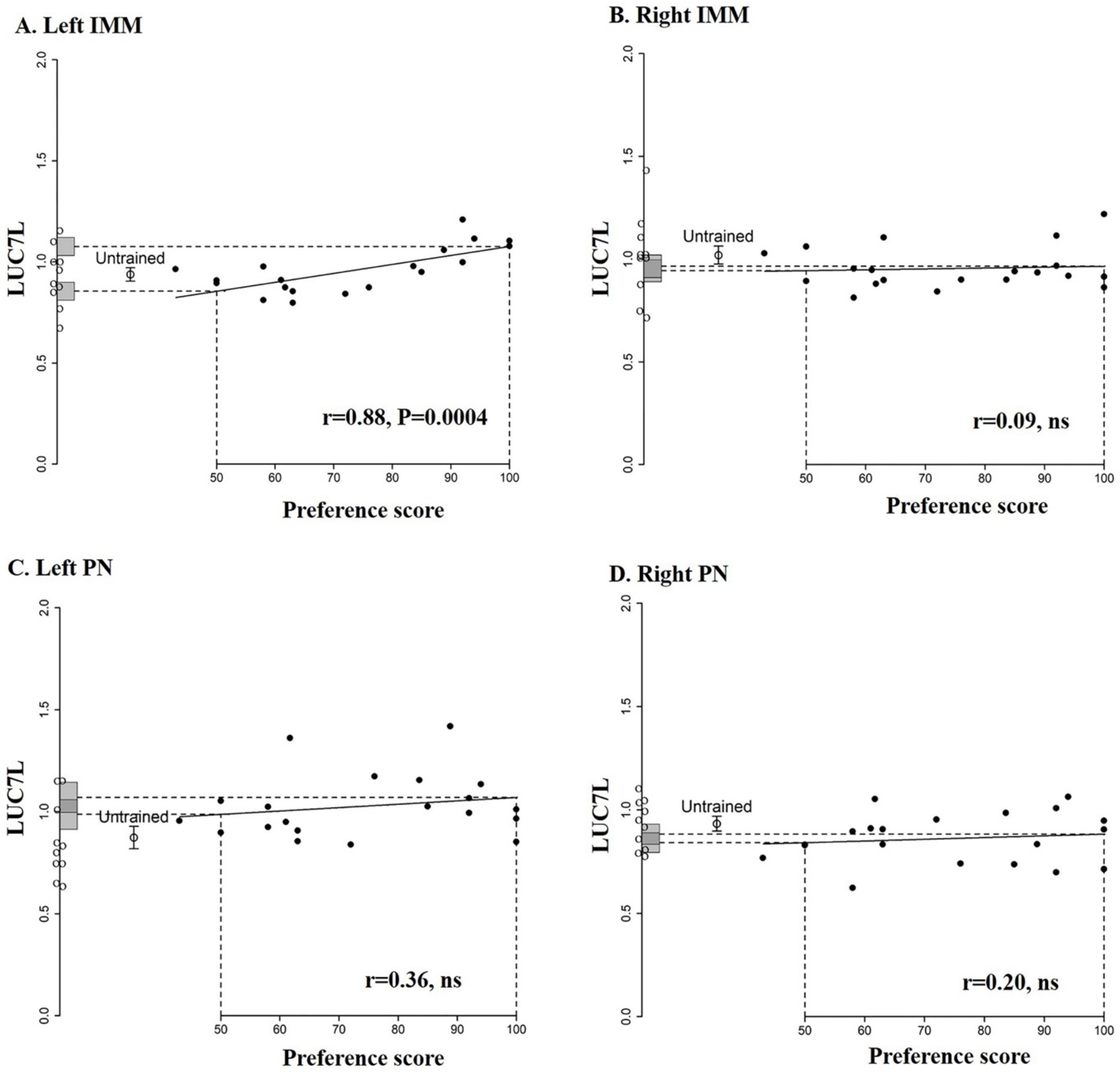

*Supplementary Figure 17: LUC7 association with memory strength, 24h after training, in the left and right IMM, as well as in the left and right PN. Standardized relative amounts of LUC7L (y-axis) are plotted against preference score (x-axis). Each filled circle stands for a trained chick, while the mean level in untrained controls ± S.E. are represented by a hollow circle with error bars. Individual values for untrained chicks are shown as open circles along the y-axis. The vertical dashed line at value 50 indicates 'no preference', meaning an equal attraction to training and alternative stimuli (absence of learning), while the vertical dashed line at value 100 ('maximum preference') indicates perfect learning. The horizontal dashed lines correspond to the y intercepts for 'no preference' and 'maximum preference' scores, with gray bars on the y-axis representing ± S.E. for these horizontal dashed lines. For the left IMM the correlation is significant (P=0.0004).*

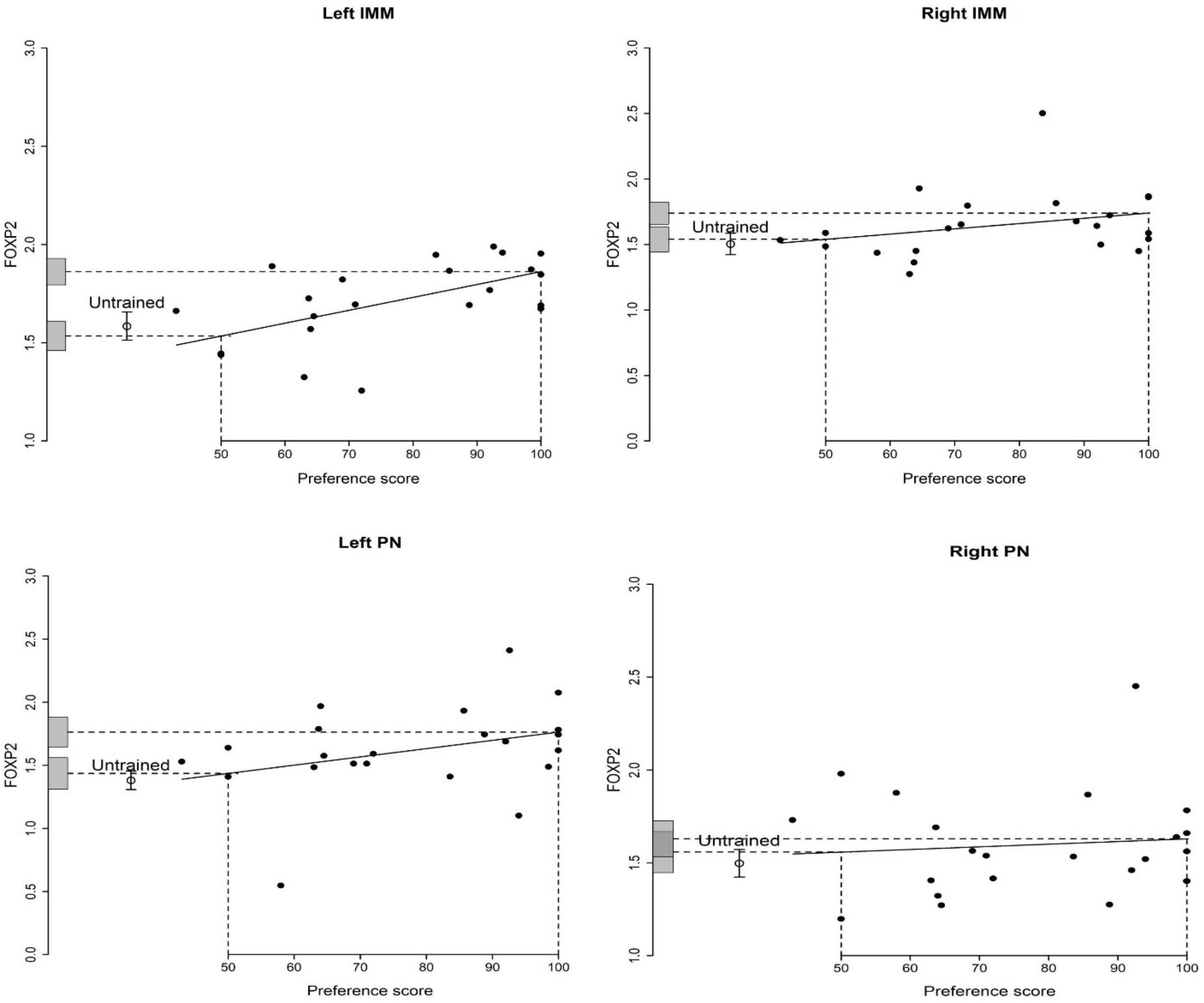

*Supplementary Figure 18: FOXP2 association with memory strength, 24h after training, in the left and right IMM, as well as in the left and right PN. Standardized relative amounts of FOXP2 (y-axis) are plotted against preference score (x-axis). Each filled circle stands for a trained chick, while the mean level in untrained controls ± S.E. are represented by a hollow circle with error bars. Individual values for untrained chicks are shown as open circles along the y-axis. The vertical dashed line at value 50 indicates 'no preference', meaning an equal attraction to training and alternative stimuli (absence of learning), while the vertical dashed line at value 100 ('maximum preference') indicates perfect learning. The horizontal dashed lines correspond to the y intercepts for 'no preference' and 'maximum preference' scores, with gray bars on the y-axis representing ± S.E. for these horizontal dashed lines. In the left IMM and in the left PN the correlations are significant (P=0.01 and P=0.03 respectively).*

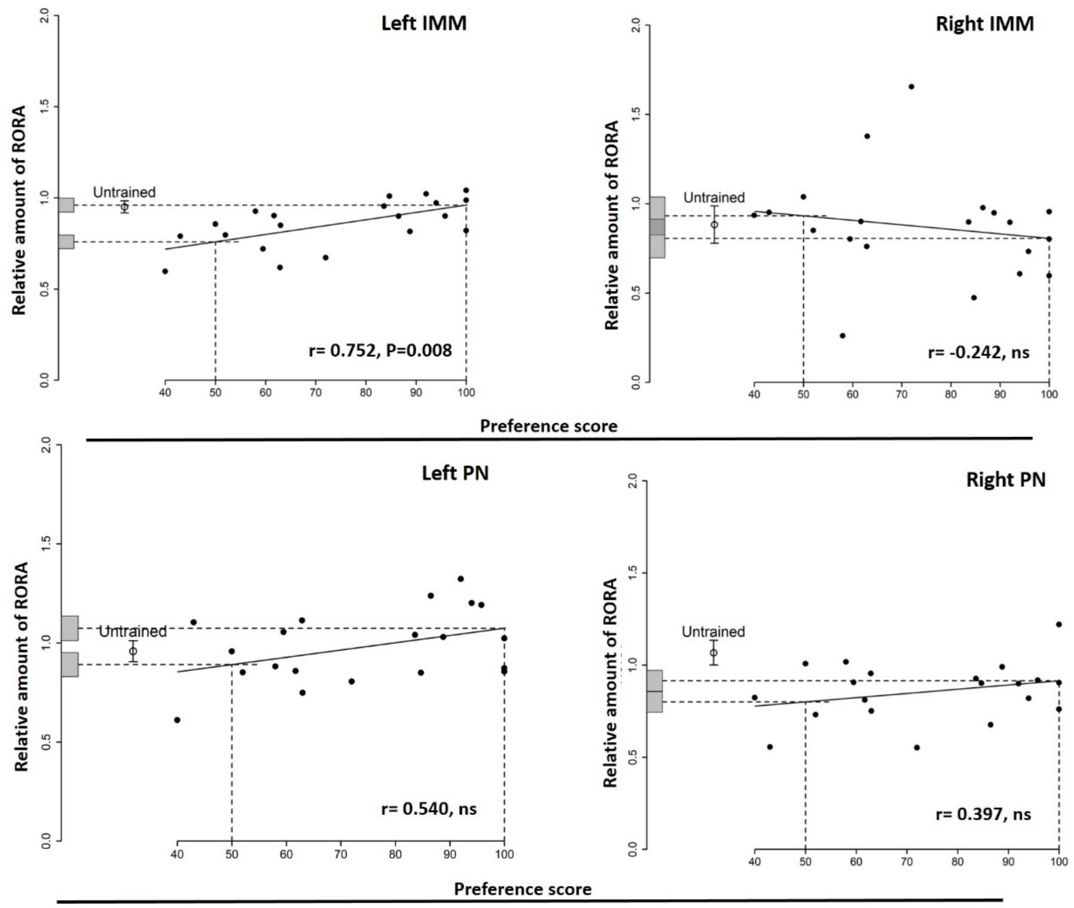

*Supplementary Figure 19: RORA association with memory strength, 24h after training, in the left and right IMM, as well as in the left and right PN. Standardized relative amounts of RORA (y-axis) are plotted against preference score (x-axis). Each filled circle stands for a trained chick, while the mean level in untrained controls ± S.E. are represented by a hollow circle with error bars. Individual values for untrained chicks are shown as open circles along the y-axis. The vertical dashed line at value 50 indicates 'no preference', meaning an equal attraction to training and alternative stimuli (absence of learning), while the vertical dashed line at value 100 ('maximum preference') indicates perfect learning. The horizontal dashed lines correspond to the y intercepts for 'no preference' and 'maximum preference' scores, with gray bars on the y-axis representing ± S.E. for these horizontal dashed lines. In the left IMM the correlation is significant (P=0.007).*

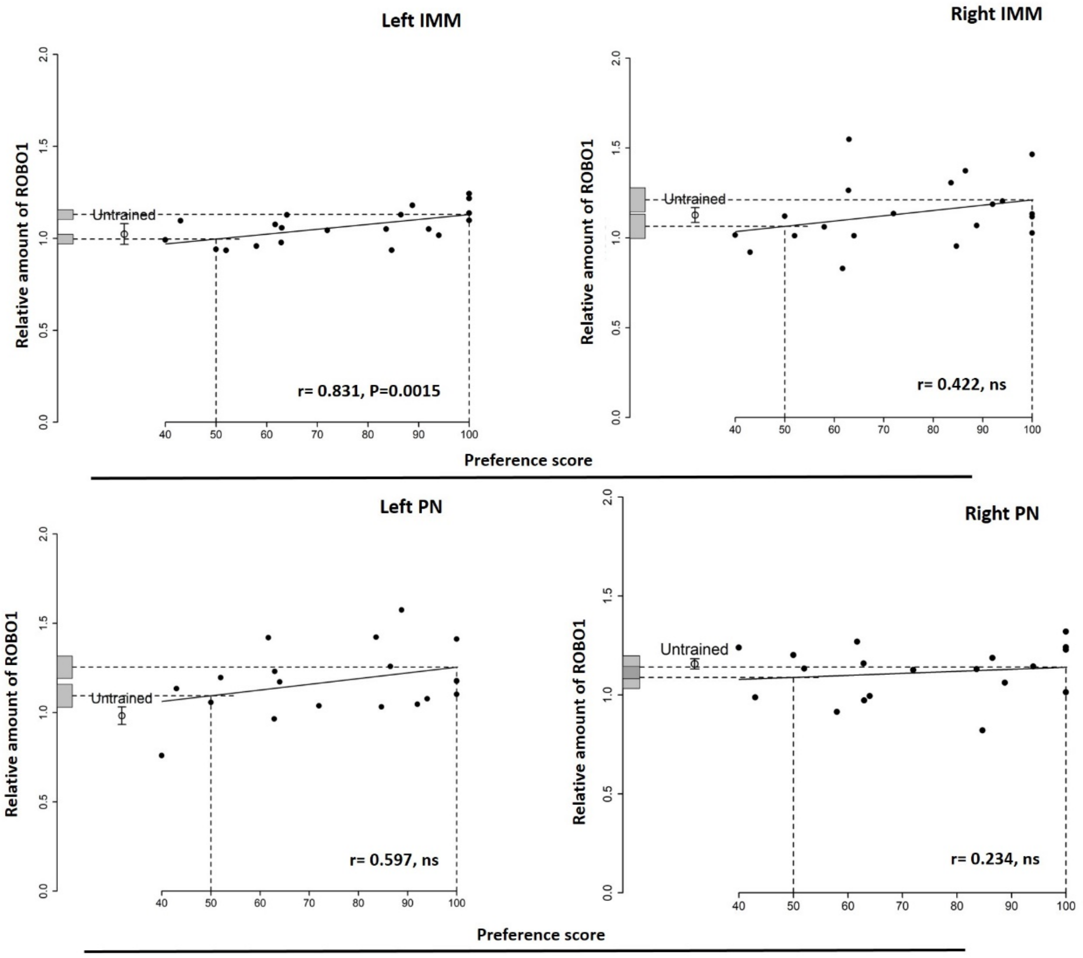

*Supplementary Figure 20: ROBO1 association with memory strength, 24h after training, in the left and right IMM, as well as in the left and right PN Standardized relative amounts of ROBO1 (y-axis) are plotted against preference score (x-axis). Each filled circle stands for a trained chick, while the mean level in untrained controls ± S.E. are represented by a hollow circle with error bars. Individual values for untrained chicks are shown as open circles along the y-axis. The vertical dashed line at value 50 indicates 'no preference', meaning an equal attraction to training and alternative stimuli (absence of learning), while the vertical dashed line at value 100 ('maximum preference') indicates perfect learning. The horizontal dashed lines correspond to the y intercepts for 'no preference' and 'maximum preference' scores, with gray bars on the y-axis representing ± S.E. for these horizontal dashed lines. In the left IMM the correlation is significant (P=0.002). Residual variance from the regression line being significantly lower than the variance of untrained chicks (P< 0.0005)*

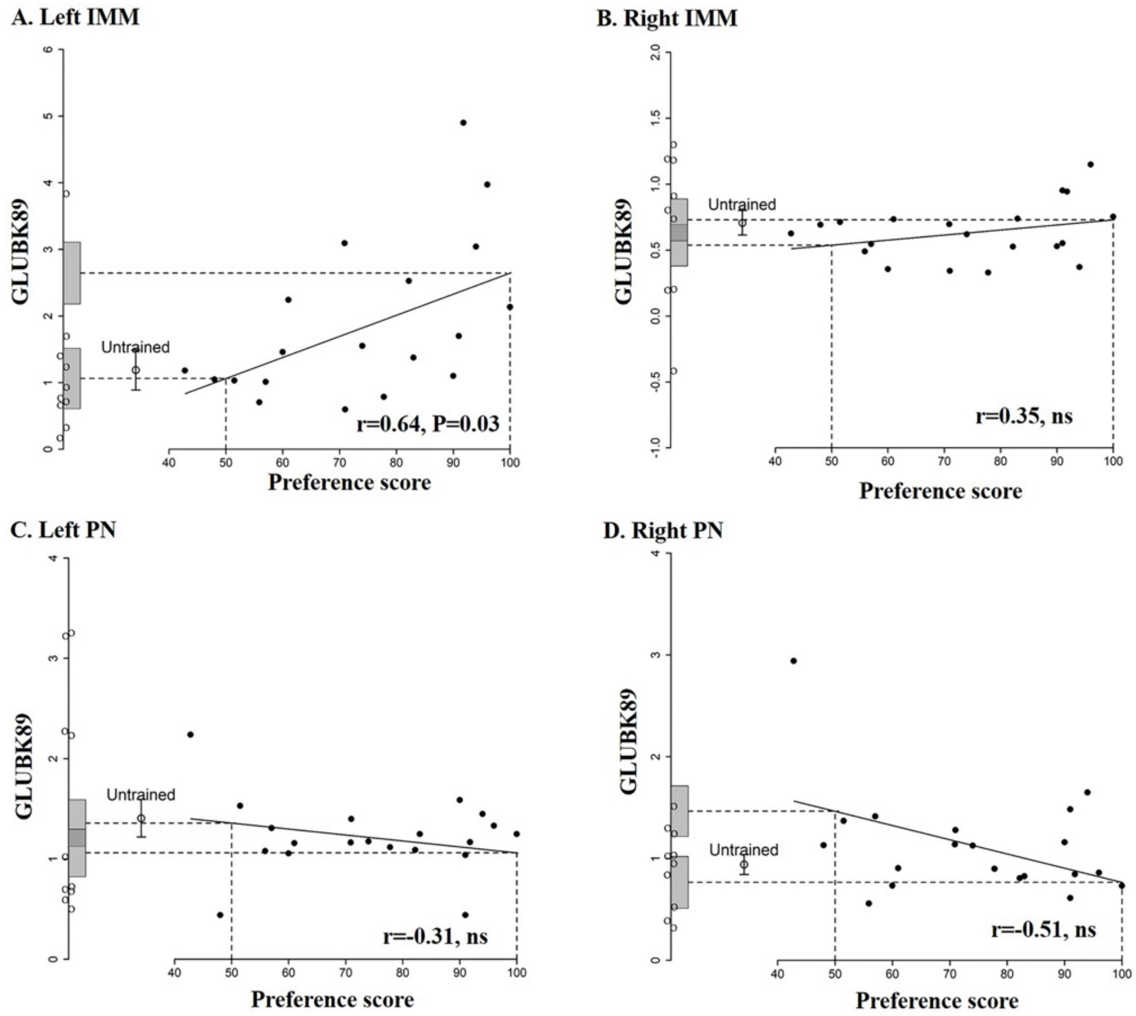

*Supplementary Figure 21: GLUBK89 association with memory strength 24h after training in the left and right IMM, as well as in the left and right PN. Standardized relative amounts of GLUBK89 (y-axis) are plotted against preference score (x-axis). Each filled circle stands for a trained chick, while the mean level in untrained controls ± S.E. are represented by a hollow circle with error bars. Individual values for untrained chicks are shown as open circles along the y-axis. The vertical dashed line at value 50 indicates 'no preference', meaning an equal attraction to training and alternative stimuli (absence of learning), while the vertical dashed line at value 100 ('maximum preference') indicates perfect learning. The horizontal dashed lines correspond to the y intercepts for 'no preference' and 'maximum preference' scores, with gray bars on the y-axis representing ± S.E. for these horizontal dashed lines. In the left IMM the correlation is significant (P=0.03). The intercept at maximum preference is significantly higher than the untrained mean (p-value = 0.019), whereas the intercept value at 50% (no learning) is not significantly different from the mean untrained value.*

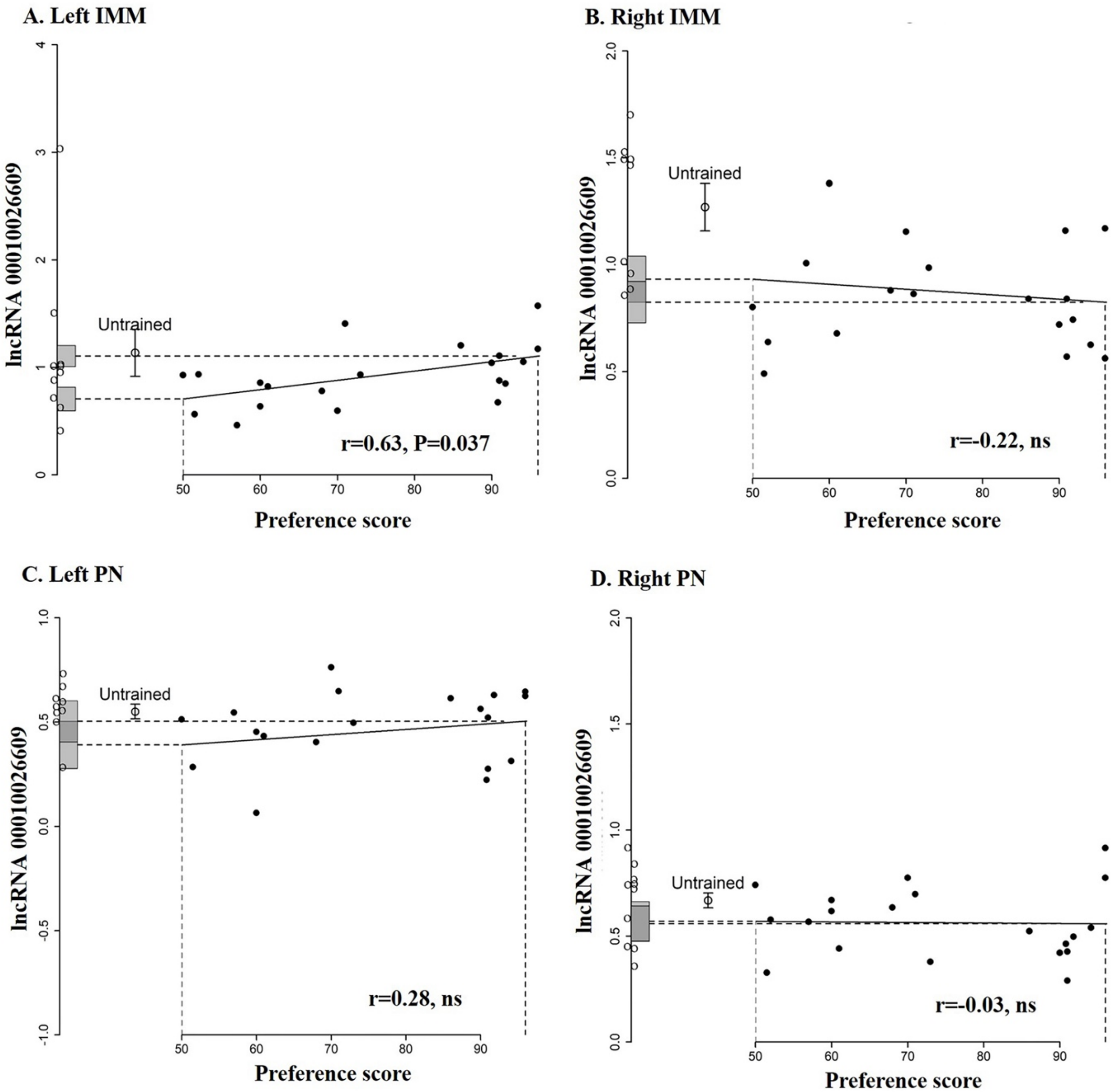

*Supplementary Figure 22: ENSGALG00010026609 lncRNA association with memory strength, 24h after training, in the left and right IMM, as well as in the left and right PN. For the left IMM the correlation is significant. Standardized relative amounts of ENSGALG00010026609 (y-axis) are plotted against preference score (x-axis). Each filled circle stand for a trained chicks, while the mean level in untrained controls ± S.E. are represented by an hollow circle with error bars. Individual values for untrained chicks are shown as open circles along the y-axis. The vertical dashed line at value 50 indicates 'no preference', meaning an equal attraction to training and alternative stimuli (absence of learning), while the vertical dashed line at value 100 ('maximum preference') indicates perfect learning. The horizontal dashed lines correspond to the y intercepts for 'no preference' and 'maximum preference' scores, with gray bars on the y-axis representing ± S.E. for these horizontal dashed lines. In the left IMM the correlation is significant (P=0.037). Residual variance from the regression line being significantly lower than the variance of untrained chicks (p-value = 0.003)*

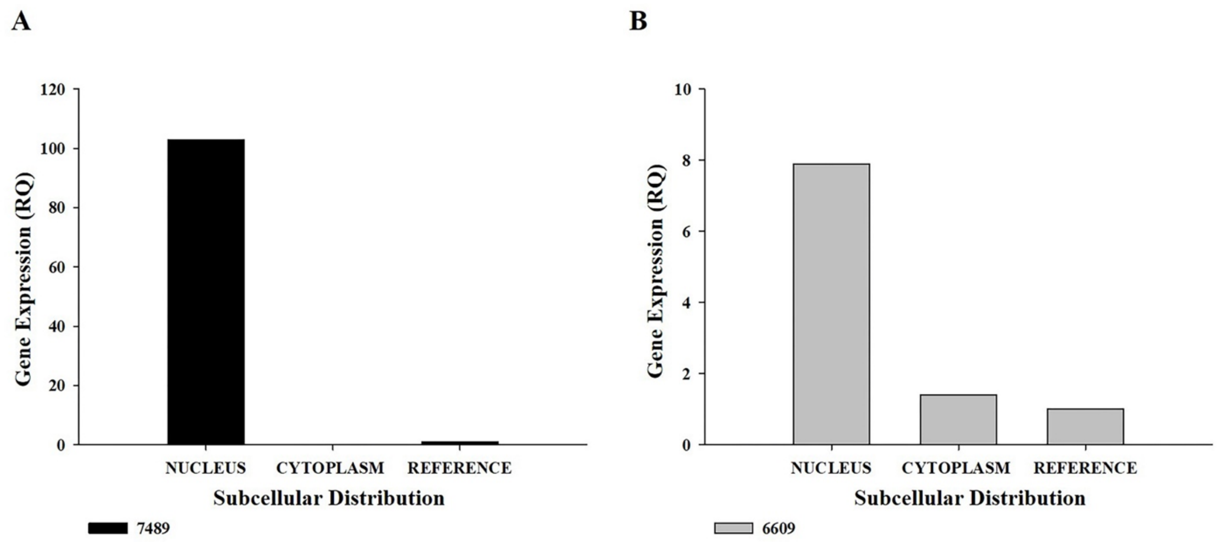

*Supplementary Figure 23: distribution of GLUBK89 (Panel A) and LNCRNA6609 (Panel B) lncRNAs in nucleus and cytoplasm.*

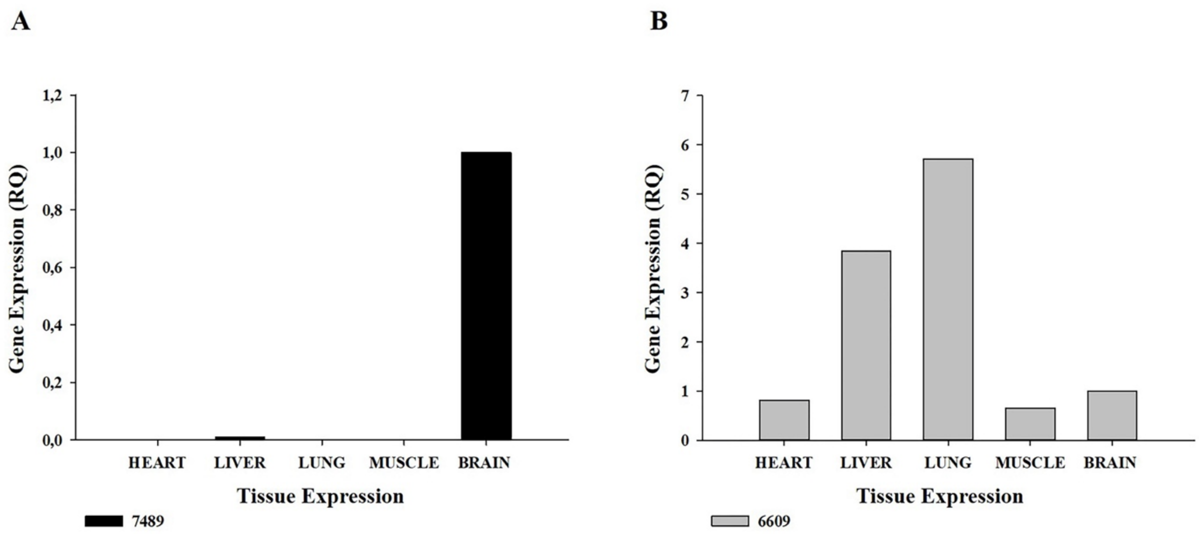

*Supplementary Figure 24: distribution of GLUBK89 (Panel A) and LNCRNA6609 (Panel B) lncRNAs across different tissues.*

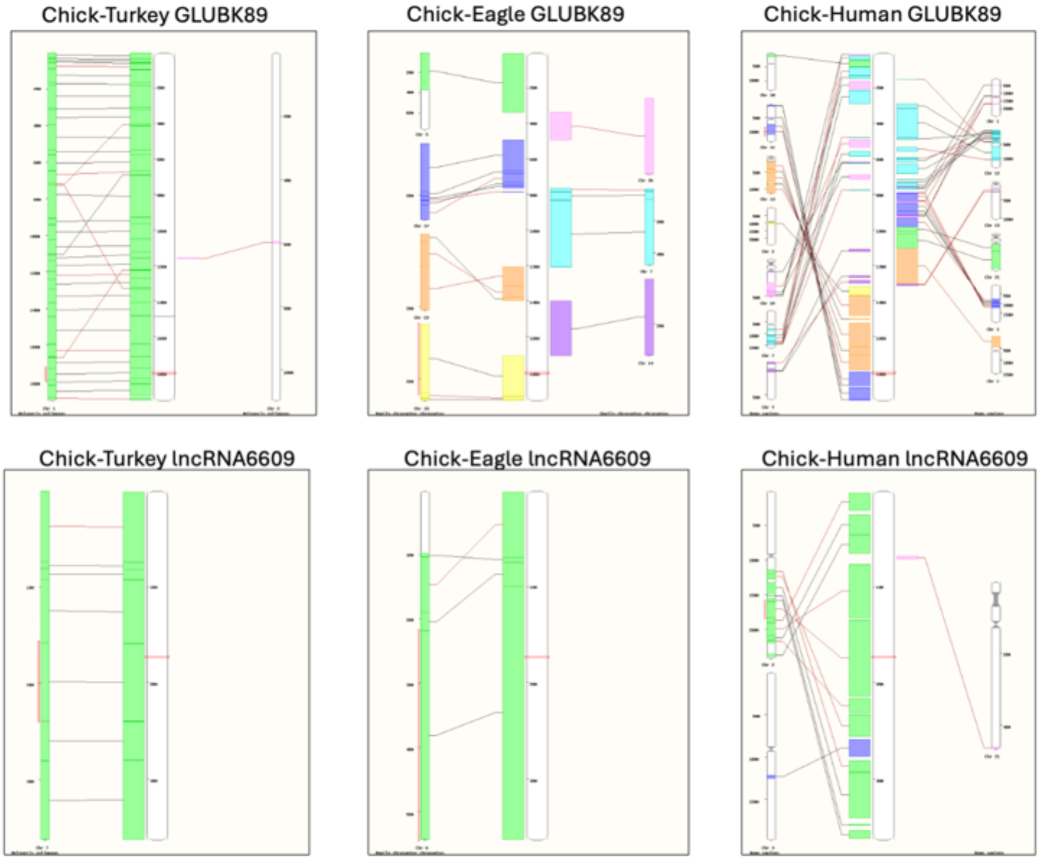

*Supplementary Figure 25: Chromosomal-level homology analyses of the chicken GLUBK89 and lncRNA6609 loci with turkey, golden eagle, and human genomes were conducted using the Ensembl comparative genomics (synteny) tool (see text for details). Syntenic regions are calculated from pairwise (two-species) whole genome alignments. The center chromosome represents the species of interest, and the smaller chromosomes show syntenic regions with a second species. Blocks are colored according to the chromosome number on the second species. Black lines connect syntenic blocks with the same orientation. Brown lines indicate regions with opposite orientation. The small red boxes mark the gene of interest and its homologue.*

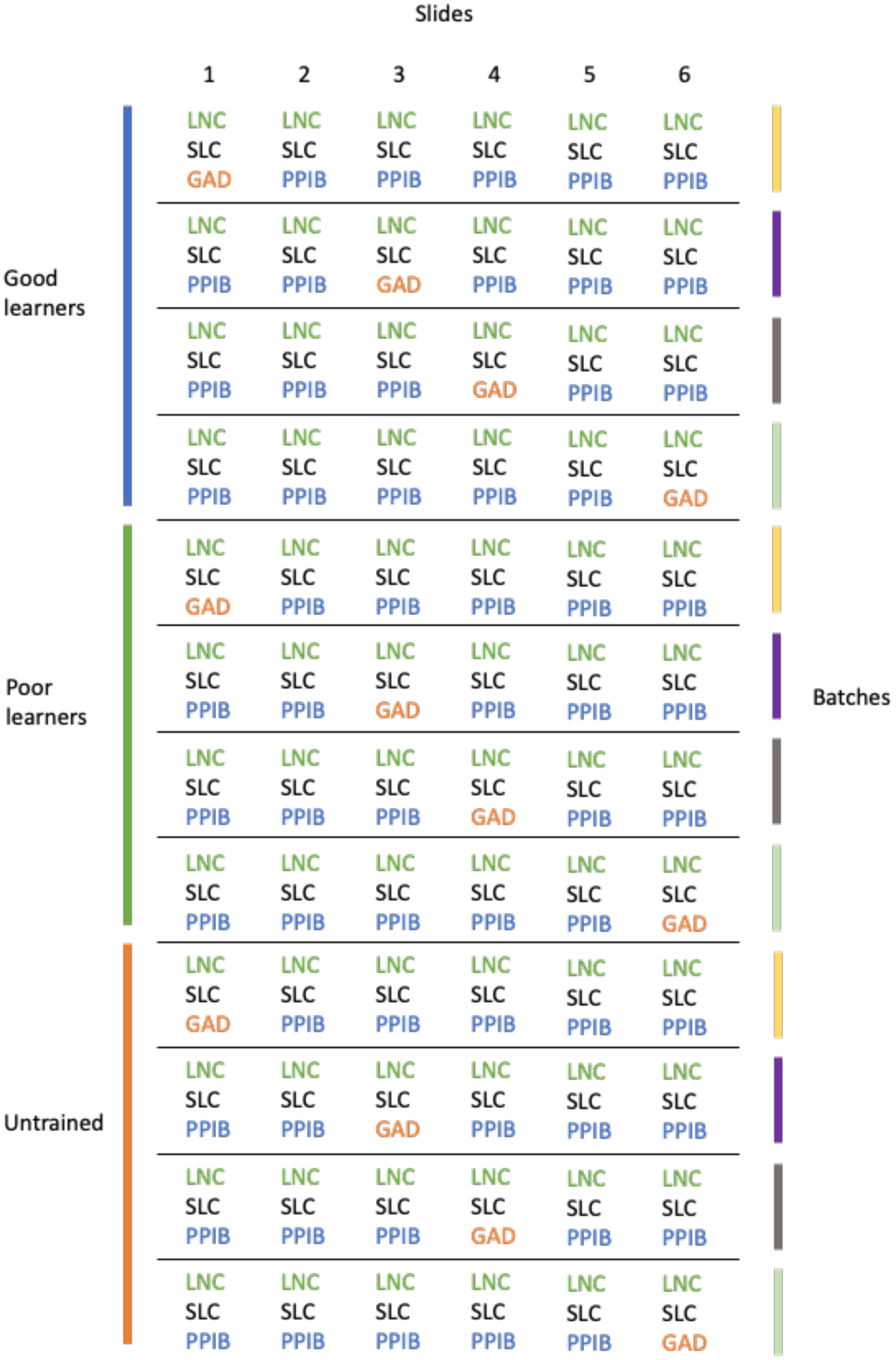

*Supplementary Figure 26: design of the in-situ experiment. Each row of the matrix represents one chick brain (sample), each column a section. Samples are grouped in three experimental groups (Good Learners, Poor Learners, Untrained, color bars on the left side), and were processed in four different batches, represented by the colored bars on the right of the figure. Five sections for each brain were stained for the long non-coding RNA ENSGALG00010007489 (LNC), the glutamatergic neurons marker SLC17A6 (SLC), and the housekeeping gene PPIB, while one section had the housekeeping gene swapped with the GABAergic neurons marker GAD2 (GAD).*

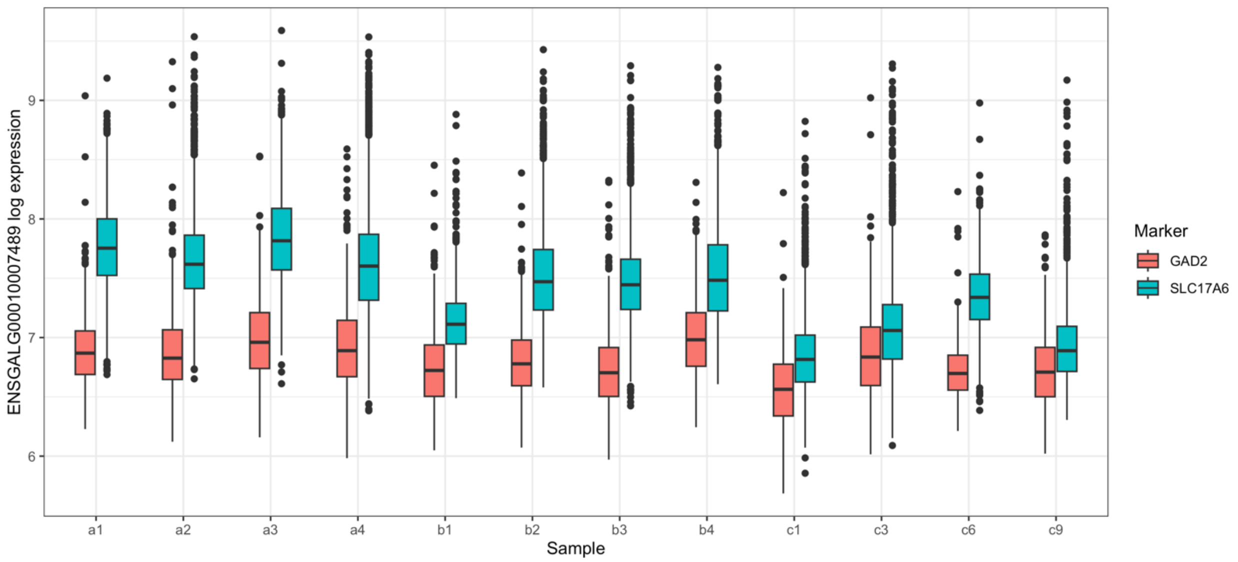

*Supplementary Figure 27: expression levels of the ENSGALG00010007489 lncRNA in chick brain nuclei, as quantified by the QuPath software on in-situ hybridized brain sections. The x-axis reports the chick (sample) from which the section was extracted. Samples' names also indicated the experimental group, namely Good Learners (names starting with a), Poor Learners (with b), and untrained (with c). Colors indicate whether the nuclei present a high level of GABAergic marker GAD2 or glutamatergic marker SLC17A6. For all samples, glutamatergic neurons have higher levels of ENSGALG00010007489 expression, in a statistically significant way (two-tailed t-test p-value < 0.001).*

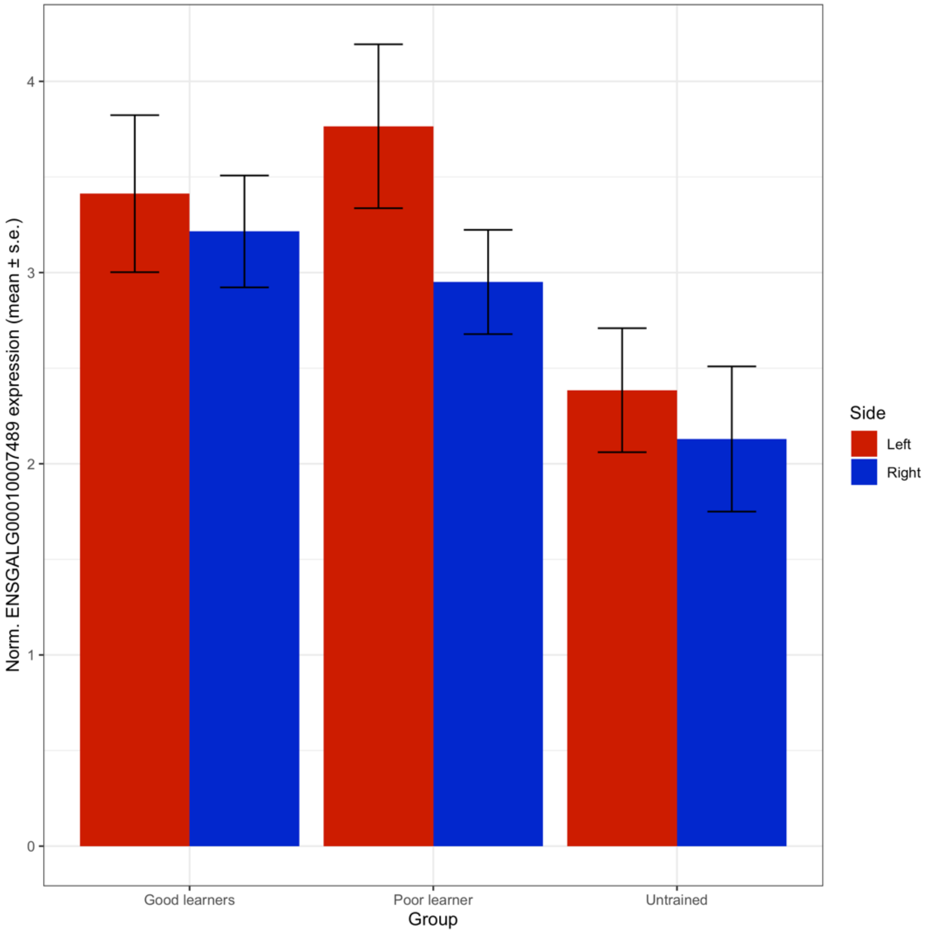

*Supplementary Figure 28: GLUBK89 expression (normalized by PPIB) in the nidopallium across groups and nidopallium sides (left, right). Height of the bars represent average value, whiskers ± standard error (s.e.).*

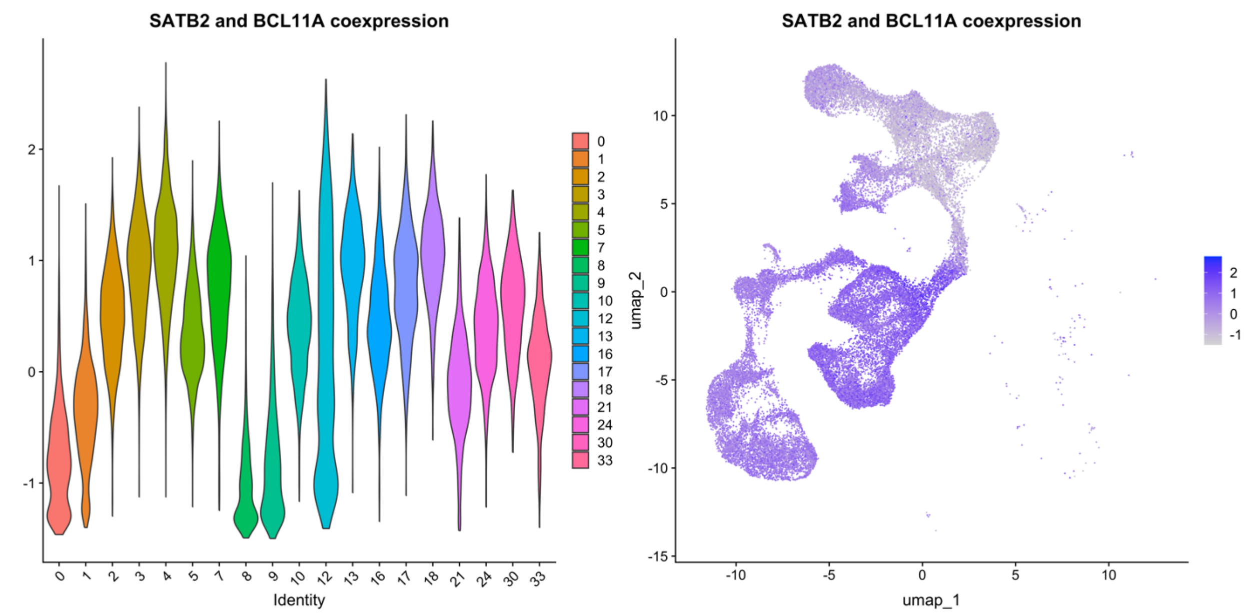

*Supplementary Figure 29: co-expression of the inter telencephalic (IT) glutamatergic markers SATB2 and BCL11A. A single module score for both genes was computed with the Seurat function "AddModuleScore"*

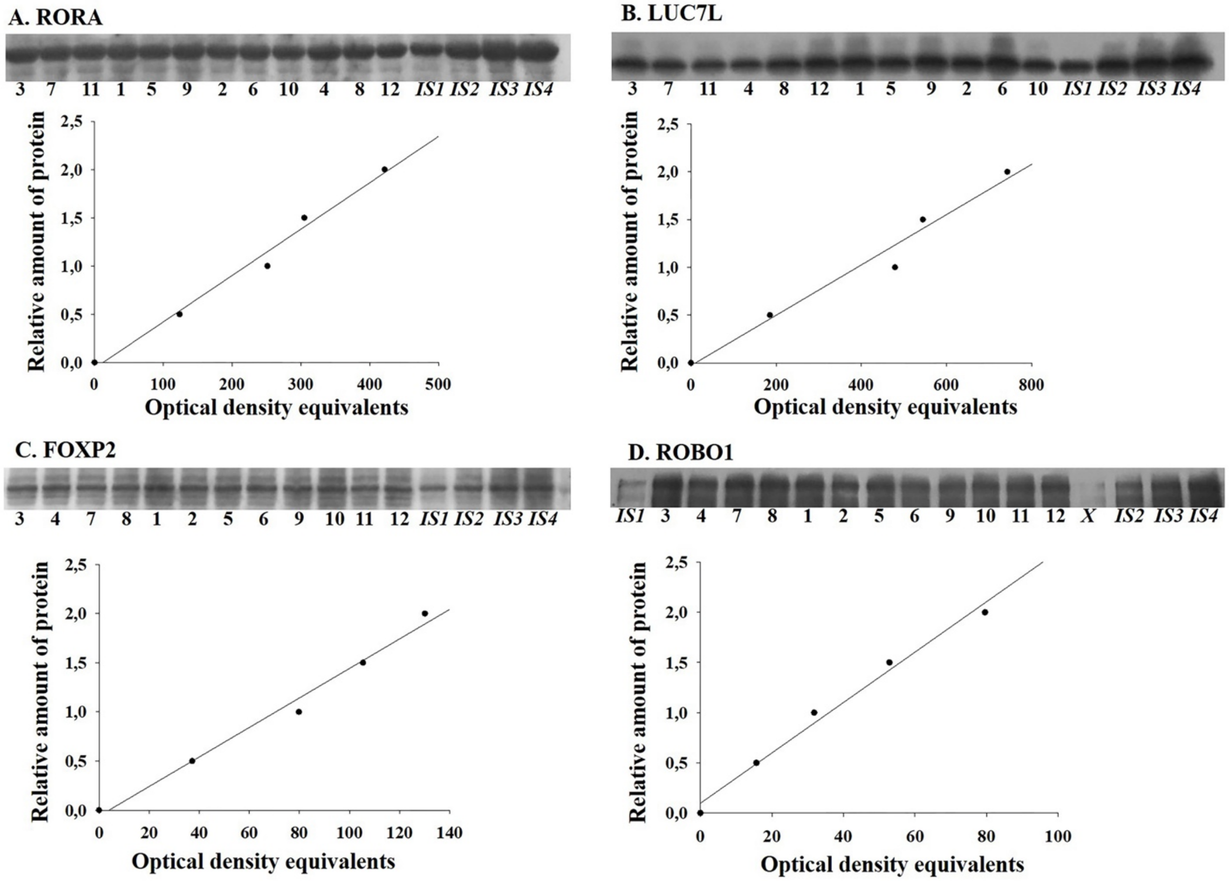

*Supplementary Figure 30: immunoblotting results. Each panel (A, B, C, D) reports the results for a single protein (RORA, LUC7L, FOXP2, ROBO1, respectively). The top of each panel reports the stained protein bands; the bottom presents the optimal density equivalents against the relative amount of each protein.*

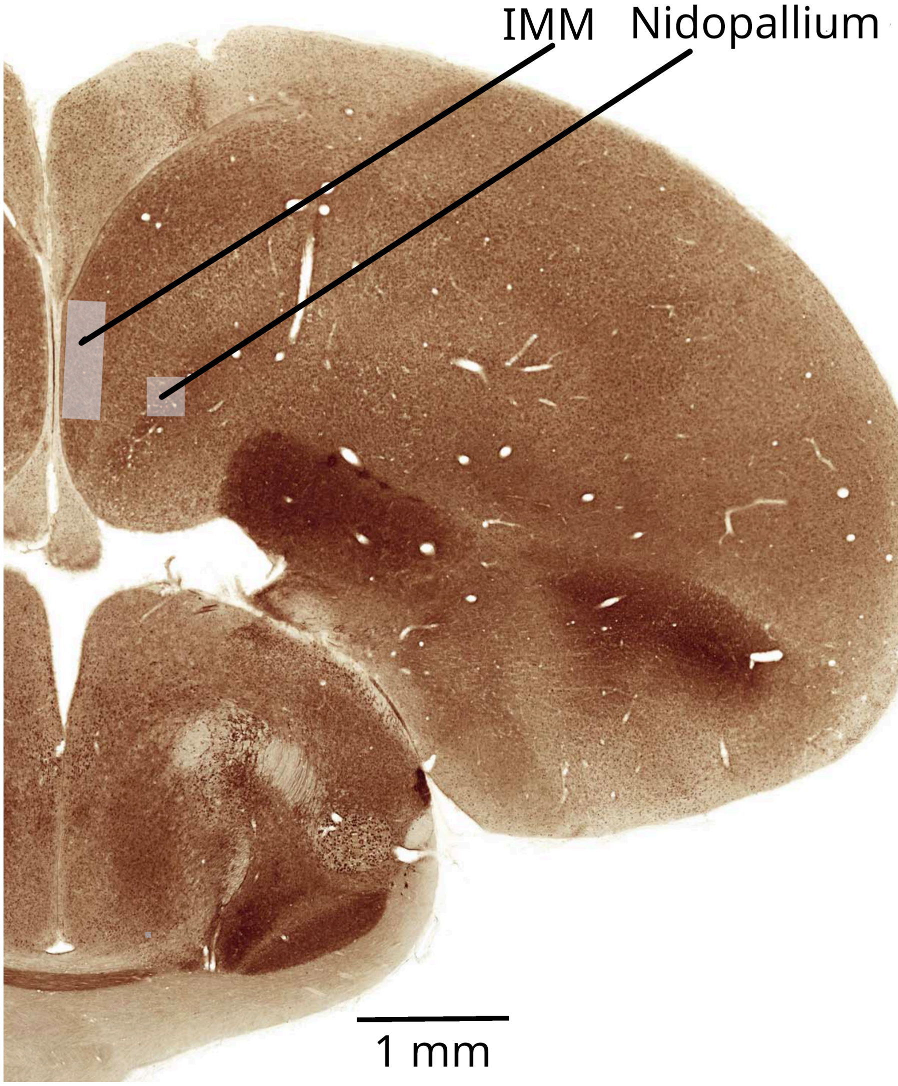

*Supplementary Figure 31: Sampling regions in the IMM and nidopallium, shown as pale rectangles superimposed on a section of chick brain stained for acetylcholinesterase. The figure is modified from Plate 17 of an atlas of the 14-day post-hatch chick brain[1]. Linear measurements have been scaled by 0.75 to give representative distances in the brain of a chick 1-2 days after hatching*
